# Three positively charged binding sites on the eastern equine encephalitis virus E2 glycoprotein coordinate heparan sulfate- and protein receptor-dependent infection

**DOI:** 10.1101/2024.11.04.621500

**Authors:** Maria D.H. Alcorn, Chengqun Sun, Theron C. Gilliland, Tetyana Lukash, Christine M. Crasto, Saravanan Raju, Michael S. Diamond, Scott C. Weaver, William B. Klimstra

## Abstract

Naturally circulating strains of eastern equine encephalitis virus (EEEV) bind heparan sulfate (HS) receptors and this interaction has been linked to its neurovirulence. Previous studies associated EEEV-HS interactions with three positively charged amino acid clusters on the E2 glycoprotein. One of these sites has recently been reported to be critical for binding EEEV to very-low-density lipoprotein receptor (VLDLR), an EEEV receptor protein. The proteins apolipoprotein E receptor 2 (ApoER2) isoforms 1 and 2, and LDLR have also been shown to function as EEEV receptors. Herein, we investigate the individual contribution of each HS interaction site to EEEV HS- and protein receptor-dependent infection *in vitro* and EEEV replication in animals. We show that each site contributes to both EEEV-HS and EEEV-protein receptor interactions, providing evidence that altering these interactions can affect disease in mice and eliminate mosquito infectivity. Thus, multiple HS-binding sites exist in EEEV E2, and these sites overlap functionally with protein receptor interaction sites, with each type of interaction contributing to tissue infectivity and disease phenotypes.

## INTRODUCTION

Eastern equine encephalitis virus (EEEV) is a member of the *Alphavirus* genus, which is comprised mostly of mosquito-borne positive-sense single-stranded RNA viruses in the family *Togaviridae*^1^. EEEV is endemic in North America. EEEV disease in humans is uniquely neurovirulent among alphaviruses, marked by limited febrile prodrome and rapid onset of encephalitis with a case fatality rate of 30-70% and a high rate of long-term neurological sequelae^1^. Currently, there are no licensed vaccines or therapeutics to prevent or treat human EEEV infections. Similar to humans, mice infected with EEEV show limited signs of prodrome, often first showing signs of encephalitis before rapidly succumbing to disease^2-4^. Consistent with the disease signs, EEEV infection exhibits limited replication in lymphoid tissues, does not induce a robust systemic innate immune response, and replication is poorly controlled in central nervous system (CNS) tissues^3,4^. These phenotypes have been linked, in part, to the ability of wild-type (WT) EEEV to efficiently bind heparan sulfate (HS) proteoglycan attachment receptors on cells^3-6^.

To date, EEEV is the only arbovirus for which amino acid residues in attachment proteins of naturally circulating, unpassaged, viruses have been shown to promote efficient binding to HS^3,7^. Previous infection-based studies identified positively charged lysine (K) residues (E2 K71, K74, and K77) in a cluster in domain A of the EEEV E2 glycoprotein as critical for HS-dependent infection^3,4^. Subsequently, cryo-electron microscopy (cryo-EM) identified two sites involving lysine (K) and arginine (R) residues in the EEEV E2 that bind a low-molecular-weight HS analog, 6kD heparin, as putative HS-binding sites (E2 R84 and R119 coordinating with HS in one site and K156, R157 in another)^8^. However, it is currently unclear whether the heparin-binding sites identified by cryo-EM contribute to HS-dependent infection or replication of EEEV in vertebrates or mosquito vectors.

More recently, several low-density lipoprotein (LDL) binding proteins, including very-low-density lipoprotein receptor (VLDLR), LDLR, and apolipoprotein E receptor 2 (ApoER2), were identified as EEEV attachment and entry receptors^9,10^. Unexpectedly, the positively charged residues in the K156 and R157 HS-binding site were among the VLDLR-E2 contact residues mapped by structural analysis^11-13^. It has been hypothesized that infectivity increases promoted by HS proteoglycans result from concentration of virus particles on cell surfaces such that interaction with *bona fide* “entry receptors” is non-specifically enhanced by HS^7,14-16^. However, the relationship of HS proteoglycan attachment receptors to protein attachment and entry receptors is unknown for RNA viruses.

Using charged-to-alanine (A) mutagenesis, we show that each of the three HS-binding sites on E2 contributes to EEEV-HS interactions at the cell surface, although to different degrees; that each site may interact with HS in a functionally unique manner; and that charged-to-alanine mutation at each of the three HS-binding sites decreases neurovirulence in mice and mosquito infectivity. Surprisingly, mutagenesis of each HS-binding site either diminished or abrogated interactions with VLDLR, ApoER2, and LDLR receptors as measured by infectivity enhancement and binding to receptor-overexpressing cells, or competition with a VLDLR-derived infection inhibitor both *in vitro* and *in vivo*. Thus, the sites present in naturally circulating EEEV glycoproteins are polyfunctional, coordinating interactions with both HS and protein receptors. Notably, abrogation of protein receptor interactions with a 156-157 (E2 K156A and R157A) mutant eliminated mosquito infection but had only a modest effect on virulence in mice, suggesting the presence of additional entry mediating factors in mammals. Finally, mutations that enhance both protein receptor- and HS-dependent infectivity could be selected through *in vitro* passage of the attenuated chimeric mutant EEEV viruses. Together, these data suggest structural and/or functional convergence between EEEV engagement with HS and protein receptors and provide a potential mechanism for historical observations of *in vitro* selection of single-site substitution mutations that confer HS-binding^17,18^.

## RESULTS

### Mutations to each HS interaction site of E2 independently alter EEEV-HS interactions

Previously, residues of EEEV E2 (K71, K74, and K77) were shown to be critical for EEEV-HS interactions, and a charge-to-alanine mutant, referred to as EEEV 71-77 ablated EEEV-HS interactions^3,4^. We, with collaborators, recently demonstrated that residues in two other sites of E2 (R84 and R119 or K156 and R157) interact with 6kD heparin^8^. To determine the extent that residues in these newly identified E2 sites contribute to EEEV-HS interactions and how they compare to the 71-77 mutant, we tested the infection characteristics of charged-to-alanine mutants at each HS/heparin-binding site with a series of HS infectivity dependence assays.

Using chimeric viruses derived from a cDNA clone encoding Sindbis virus (SINV) nonstructural proteins and RNA replication control structures^19,20^, and expressing EEEV structural proteins (SINV/EEEV), we compared the infectivity of WT and each mutant on CHO-K1 cells *versus* glycosaminoglycan (GAG)-deficient CHO-pgsA-745 and HS-deficient CHO-pgsD-677 cells^21^. Consistent with previous results^3,4^, SINV/EEEV WT infectivity was significantly reduced (p<0.001) in the absence of all GAGs or HS, whereas the 71-77 mutant was unaffected. The 84-119 (E2 R84A, R119A) mutant exhibited reduced infectivity similar to the WT, whereas the 156-157 mutant was similar to 71-77, showing no significant reduction in infectivity for the mutant cells (**Fig. 1A**). GAG-dependent infectivity was recapitulated in cellular binding assays, with WT and the 84-119 mutant showing a significant decrease in binding to CHO-psgA-745 compared to CHO-K1 cells, and the 71-77 and 156-157 mutants showing decreased binding to CHO-K1 cells compared to the WT virus and similar binding to both WT and mutant cell lines (**Fig. 1B**). We then compared the infection rates of each mutant when incubated in media with increased NaCl concentration, which disrupts ionic interactions such as protein-HS binding^22-25^. Consistent with previous data^4^, the WT showed a significant decrease in infectivity starting at the lowest concentration of supplemented NaCl, whereas the 71-77 mutant was not susceptible to ionic disruption (**Supp. Fig. 1A**). The 84-119 and the 156-157 mutants exhibited intermediate phenotypes, showing significantly reduced infectivity compared to an RPMI control but higher infectivity compared to WT at the higher NaCl concentrations (**Supp. Fig. 1A**).

**Fig. 1.**
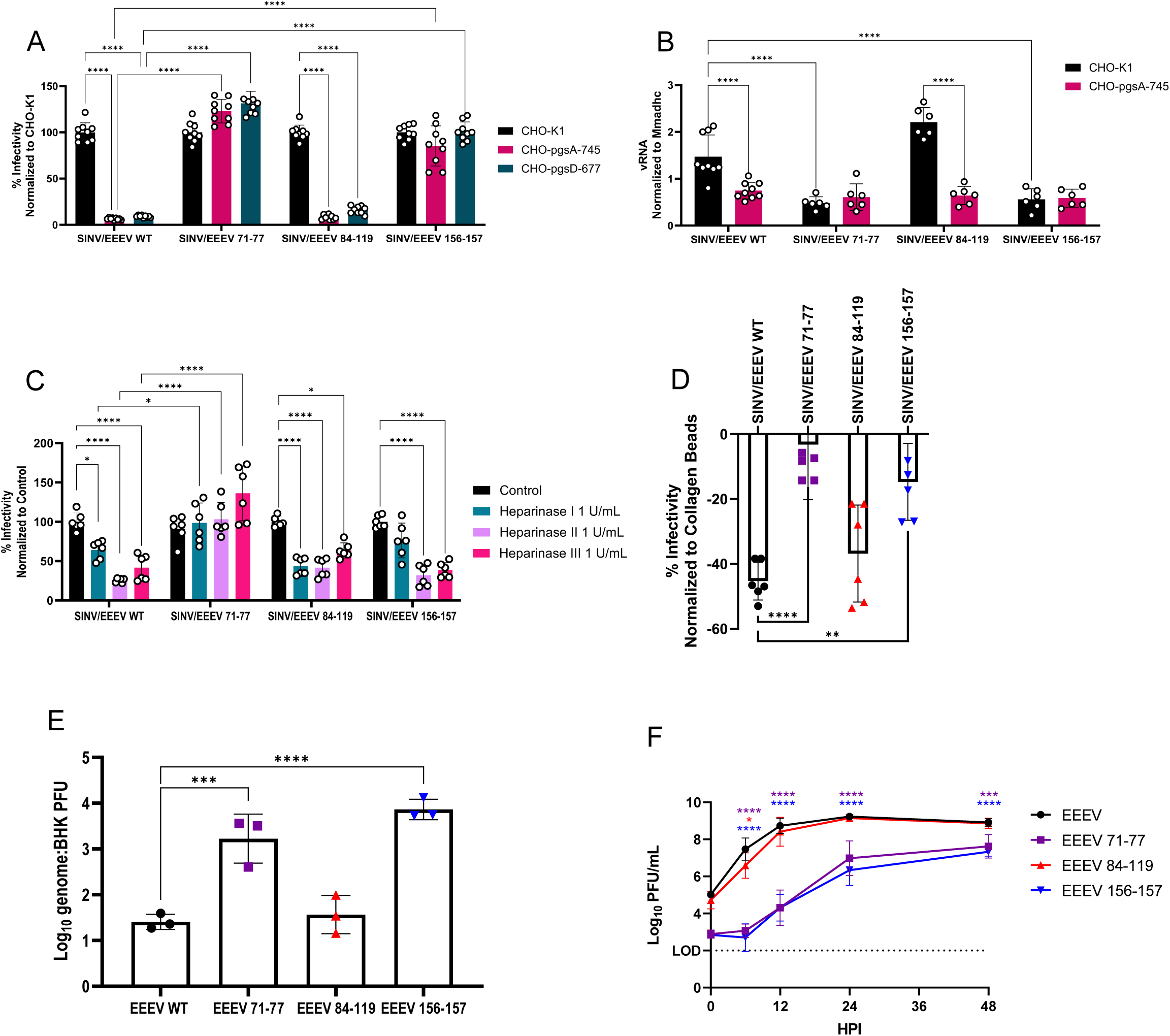
E2 mutations independently and uniquely alter EEEV-HS interactions *in vitro*. **A** Relative infectivity of WT or mutant SINV/EEEV for CHO-K1, GAG-deficient CHO-pgsA-745, and HS-deficient CHO-pgsD-677 cells (n = 9 from 3 independent experiments). Viruses were titrated on all three cell types, and percent infectivity compared to CHO-K1 was determined for each cell type. **B** Binding of chimeric WT and mutant viruses to CHO-K1 and CHO-pgsA-745 cells (n = 6 from 2 independent experiments, except for SINV/EEEV WT where n = 9 from 3 independent experiments). Viruses were allowed to incubate with cells on ice for 75 minutes, then cells were washed, and the bound virus was quantified by ddPCR as a ratio of genomes bound to cells *versus* the host *Mmadhc* gene. **C** Relative infectivity of WT or mutant SINV/EEEV on BHK-21 cells treated with heparinases (n = 6 from 2 independent experiments). **D** Relative indirect binding to heparin-agarose beads by WT and mutant SINV/EEEV. Viruses were incubated with collagen-agarose or heparin-agarose beads; unbound viruses were titered on BHK-21 cells (n = 6 from 2 independent experiments). **E** Genome-to-BHK PFU ratios for EEEV WT and mutant viruses (n = 3 independent experiments). **F** BHK-21 cells were infected with equal genomes of EEEV WT and mutant EEEVs, corresponding to a multiplicity of infection of 1 for WT (n = 6 from 2 independent experiments). All error bars show standard deviation (SD). Significance determined by (**A-C**) two-way ANOVA with Tukey’s post-hoc tests, (**D**) Brown-Forsythe and Welch ANOVA with Dunnett’s post-hoc tests, (**E**) one-way ANOVA with Tukey’s post-hoc tests on log-transformed data, or (**F**) two-way repeated measures ANOVA with Tukey’s post-hoc tests, *(p<0.05), ** (p<0.01) ***(p<0.001), ****(p<0.0001).

To determine if mutations at each site impact qualitative aspects of the EEEV interaction with cell surface HS, we examined the infectivity of chimeric WT and each mutant for BHK-21 cells following treatment with different microbial heparinases, which target different HS modifications based on sulfonation patterns^26^. The SINV/EEEV WT infectivity was significantly reduced following each heparinase treatment but was most susceptible to heparinase II or III treatments (**Fig. 1C**). In contrast, the infectivity of the 71-77 mutant was not decreased following treatment with any heparinase. The 84-119 mutant was also reduced in infectivity following treatment with each heparinase (**Fig. 1C**). However, unlike the WT, the 84-119 mutant was more significantly reduced following heparinase I or II treatments. Although the 156-157 mutant was not significantly decreased in infectivity or binding for GAG-deficient CHO cells (**Fig. 1A-B**), it was susceptible to heparinase II and III treatment similar to the WT (**Fig. 1C**). Together, CHO cell infectivity, salt disruption and heparinase digestion data suggest that each of the three mutations alter EEEV-HS interactions in a unique manner.

To determine whether the mutations impacted the ability of virions to bind to HS/heparin in the absence of other cellular receptors, we performed indirect binding assays using heparin-agarose beads (a highly sulfonated analog of HS^27^) with collagen-agarose beads serving as a control. Decreased infectivity for viruses incubated with heparin *versus* collagen demonstrates increased heparin binding. The WT bound most efficiently of the four viruses to the heparin-agarose beads, with the 84-119 mutant binding slightly, but not significantly, less. The 71-77 and 156-157 mutants exhibited significantly reduced binding to heparin beads compared to the WT and the 84-119 mutant (**Fig. 1D**).

Increased HS-binding in cell-adapted viral mutants has been associated with increased infectivity for cells and decreased genome/particle-to-plaque forming unit (PFU) ratios.^6,25,28^ We determined the genome-to-PFU ratios of each virus on BHK-21 cells. The 84-119 mutant was similar to the WT, whereas the genome-to-PFU ratios for the 71-77 and 156-157 mutants were approximately 100-fold increased compared to the WT (**Fig. 1E**), suggesting reduced infectivity per particle. To ensure that all mutants could replicate effectively, single-step replication curves of EEEV viruses were performed using BHK-21 cells (**Fig. 1F**). Overall, the 84-119 mutant replicated similarly to the WT; however, there was a significant decrease at six hours post-infection (hpi) (**Fig. 1F**). Whereas the 71-77 and 156-157 mutants replicated similarly but more slowly and to a significantly lower titer than the WT throughout the replication curve (**Fig. 1F**). However, due to differences in genome-to-PFU ratios and reduction in HS-dependent infectivity of the 71-77 and 156-157 mutants for BHK-21 cells (**Fig. 1E**), plaque titers exaggerate replication differences, which most likely reflect differences in HS interactions.

### Mutations that disrupt E2 HS binding sites also interfere with EEEV-protein receptor interactions

Previously, contact residues were mapped for chikungunya virus (CHIKV) and Venezuelan equine encephalitis virus (VEEV) interactions with the matrix remodeling associated 8 (MXRA8) and low-density lipoprotein receptor class A domain 3 (LDLRAD3) receptors, respectively^29-32^. At least one of the residues of all three EEEV HS-binding sites is either immediately adjacent to or directly overlap to contact residues for either or both receptors (**Supp. Fig. 2**). Furthermore, two recent studies mapping EEEV-VLDLR interactions, identified the E2 K156 and R157 residues as critical for engagement with the LA1 domain of VLDLR, making these residues central to a shared receptor binding motif across multiple alphaviruses^11-13^. We hypothesized that all three HS-binding mutations, especially the 156-157 mutation, may impact the ability of EEEV to interact with one or more protein receptors. We first determined whether the E2 mutations impacted SINV/EEEV infectivity for K562 cells expressing either an empty vector (EV), VLDLR or ApoER2 isoform 1 or 2^9^ (**Fig. 2A**), or THP-1 cells expressing an EV or LDLR^10^ (**Fig. 2B**). Infectivity of the 156-157 mutant and, to a lesser degree, the 71-77 mutant was significantly decreased *versus* the WT for all cells over-expressing protein receptors. Whereas the 84-119 mutant showed smaller decrease in protein-receptor mediated infectivity, exhibiting a significant decrease in protein receptor dependent-infectivity for the ApoER2 isoform 2 and LDLR only (**Fig. 2A-B**). The infectivity results were largely recapitulated when virus binding to K562-EV and K562-VLDLR cells was assessed, with the WT showing significantly higher binding to the K562-VLDLR *versus* K562-EV cells and the 84-119 mutant showing a reproducible but non-significant increase in binding to the K562-VLDLR cells. The binding of the 71-77 and 156-157 mutants was not significantly different on the two cell types and was significantly decreased with K562-VLDLR cells compared to the WT (**Fig. 2C**). While this manuscript was in preparation, another group reported that the K156A mutation disrupts VLDLR and ApoER2 binding^12^.

**Fig. 2.**
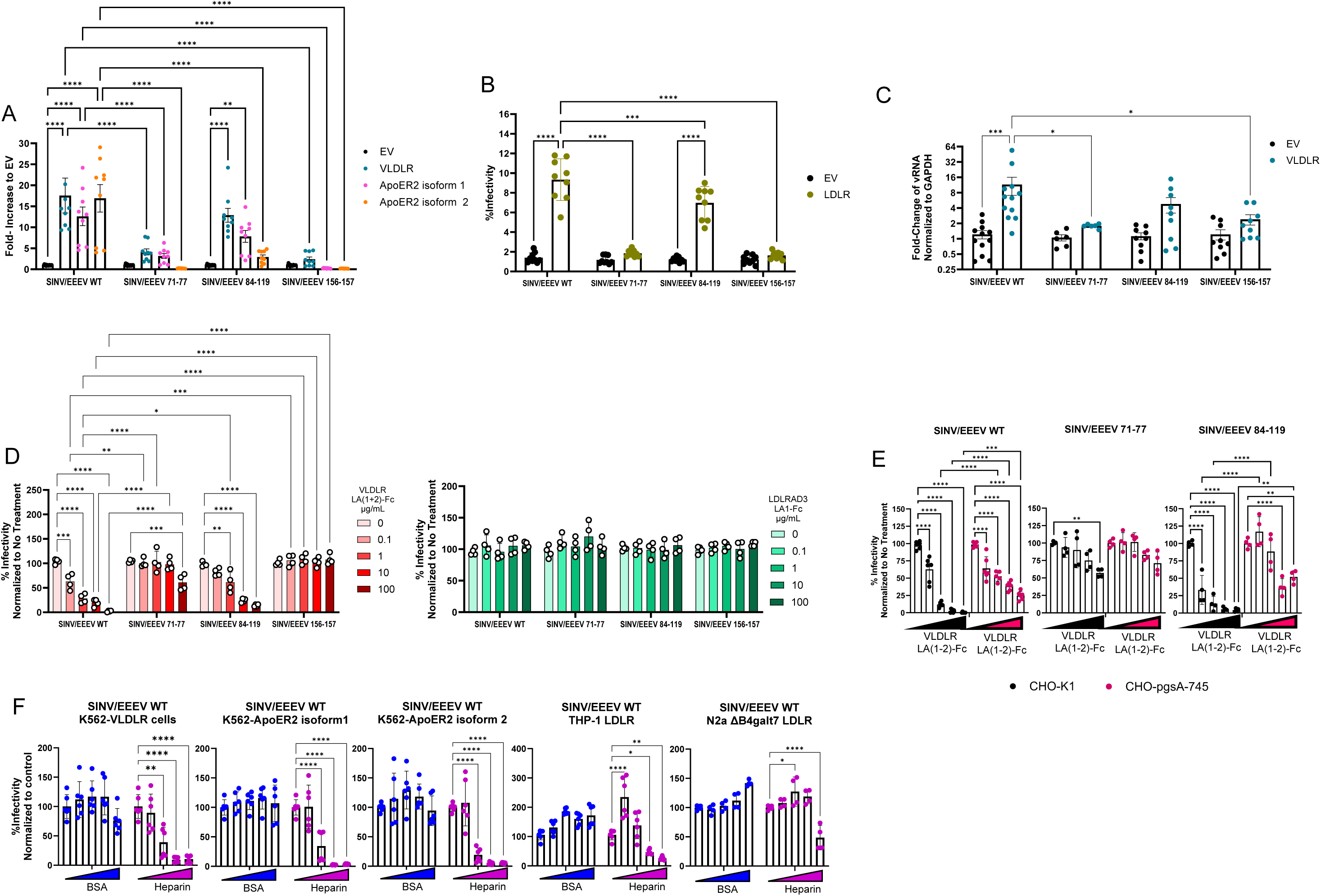
E2 mutations that disrupt E2 HS-binding sites also interfere with EEEV-protein receptor interactions. **A-B** Infection of chimeric eGFP reporter WT and mutant SINV/EEEV viruses quantified by flow cytometry. **A** K562 cells expressing empty vector (EV), VLDLR, ApoER2 isoform1, or ApoER2 isoform2; infection expressed as fold-increase (n = 9 from 3 independent experiments, except for EV where n = 15 from 5 independent experiments). **B** THP-1 cells expressing EV or LDLR; infection expressed as percent infectivity (n = 9 from 3 independent experiments). **C** Binding of chimeric WT and mutant viruses to K562-EV or K562-VLDLR cells (n = 12 from 4 independent experiments for WT, n = 9 from 3 independent experiments for 84-119 and 156-157, and n = 6 from 2 independent experiments for 71-77). Viruses were allowed to incubate with cells on ice for 75 minutes, then cells were washed, and the bound virus was quantified by qPCR, then expressed as a ratio to *GAPDH* and normalized to EV. **D** Neutralization of chimeric eGFP reporter WT and mutant viruses by VLDLR LA(1-2)-Fc or LDLRAD3 LA1-Fc (100-0.1 µg/mL in 10-fold dilutions and 0 µg control) in Vero cells (n = 4 from 2 independent experiments). **E** Neutralization of chimeric eGFP reporter WT and mutant viruses by VLDLR LA(1-2)-Fc in CHO-K1 and CHO-pgsA-756 cells (n = 4 from 2 independent experiments, except for WT where n = 9 from 3 independent experiments). **F** Neutralization of chimeric eGFP reporter WT by heparin or BSA control (2,000-2 µg/mL in 10-fold dilutions and control) in cells overexpressing protein receptors (n = 6 from 2 independent experiments) quantified by flow cytometry. Significance was determined by two-way ANOVA with Tukey’s post-hoc tests. Error bars show SD, except for (**B**) where error bars show standard error of the mean (SEM); *(p<0.05), **(p<0.01), ***(p<0.001), ****(p<0.0001).

To further assess whether or not the E2 mutations disrupted EEEV-VLDLR interactions, we performed neutralization assays using VLDLR LA(1-2)-Fc, a two-domain truncated “receptor decoy” version of the VLDLR receptor fused to the constant region of human IgG1^11^. Vero cells were infected with the SINV/EEEV WT and mutant viruses following incubation with increasing concentrations of VLDLR LA(1-2)-Fc or LDLRAD3 LA1-Fc^19^ used as a control. The WT virus was significantly neutralized at the lowest dilution of VLDLR LA(1-2)-Fc, with increased inhibition at higher concentrations (**Fig. 2D**). The 84-119 mutant exhibited a neutralization phenotype similar to the WT but was significantly more resistant to neutralization at the 1 µg/mL concentration. The 71-77 mutant was significantly less susceptible to neutralization than the WT at all concentrations and was significantly inhibited only at 100 µg/mL of VLDLR LA(1-2)-Fc (**Fig. 2D**). In comparison, the 156-157 mutant was completely resistant to neutralization by VLDLR LA(1-2)-Fc (**Fig. 2D**). Therefore, mutations at each of the three sites interfere with EEEV-VLDLR LA(1-2)-Fc interactions, with the 156-157 mutation having the greatest impact.

Given the overlap in the E2 mutations affecting HS and protein receptor interactions, we hypothesized that inhibition of EEEV infection by VLDLR LA(1-2)-Fc could partly be due to interfering with EEEV-HS interactions. To test this, we determined VLDLR LA(1-2)-Fc inhibition of SINV/EEEV WT and the 71-77 and 84-119 mutants on CHO-K1 and CHO-pgsA-745 cells. As the 156-157 mutant was not neutralized by the VLDLR LA(1-2)-Fc decoy, it was omitted in these experiments. The CHO-K1 and CHO-pgsA-745 cells both expressed similar, low levels of VLDLR but did not express ApoER2 or LDLR; thus, differences in neutralization on these cells would primarily be driven by cellular HS expression (**Supp. Fig. 3A-D**). While the WT was significantly neutralized by all concentrations of the VLDLR decoy on both cell types, at higher concentrations of inhibitor, neutralization was significantly reduced on CHO-pgsA-745 cells *versus* the CHO-K1 cells (**Fig. 2E**). The 84-119 mutant was less inhibited by the VLDLR decoy on CHO-pgsA-745 cells *versus* the CHO-K1 cells for all concentrations, only showing significant inhibition by the decoy at higher concentrations on CHO-pgsA-745 cells (**Fig. 2E**). As with Vero cells (**Fig. 2D**), the 71-77 mutant was only significantly neutralized at the highest concentration on CHO-K1 cells. In agreement with 71-77 mutant having the most profound disruption of EEEV-HS interactions, there was no difference between the neutralization of this mutant on CHO-K1 *versus* CHO-pgsA-745 cells (**Fig. 2E**). Together, these data suggest that neutralization by VLDLR LA(1-2)-Fc is aided by blockade of cellular HS-binding.

To determine whether, inversely, heparin diminishes EEEV-protein receptor interactions, we tested heparin neutralization of the SINV/EEEV WT virus on cells that over-expressed EEEV protein receptors. The WT was significantly inhibited by 20 or 200 µg/mL heparin, with increasing neutralization at higher concentrations on K562-VLDLR, K562-ApoER2, and THP-LDLR cells (**Fig. 2F**). To determine whether the high degree of neutralization was due to heparin blocking of EEEV-HS interactions and reducing overall cell attachment efficiency *versus* directly impacting protein receptor engagement, we also tested neutralization on N2a ΔB4galt7-LDLR cells, which are deficient in HS and chondroitin sulfate and over-express LDLR. On N2a ΔB4galt7-LDLR cells, heparin inhibited infection but only at the highest (2,000 µg/mL) concentration (**Fig. 2F**). Finally, treatment of K562-EV and K562-VLDLR cells with high concentrations of heparinase II demonstrated that EEEV infectivity for K562-VLDLR cells was not primarily dependent upon attachment to HS (**Supp. Fig. 1B**). Together, these data demonstrate that while heparin contributes to HS-mediated neutralization of SINV/EEEV WT on GAG+ receptor overexpressing cells, heparin does directly compete with protein receptor binding.

### Passage of HS/protein receptor binding site mutants in cultured cells selects for mutations that impact binding of both receptor types

As HS- and protein receptor-binding residues appeared to overlap functionally in that both receptors enhance infectivity, we sought to determine whether selection for partial or full restoration of HS- or protein receptor-binding would affect one or both interactions. Rapid acquisition of positively charged amino acid mutations that confer increased HS-binding after arbovirus passage in cultured cells has been extensively documented (reviewed in ^7^). We sought to mimic this process by passaging the receptor binding domain SINV/EEEV mutants on BHK-21 cells, which abundantly express HS but have low expression of VLDLR, ApoER2, or LDLR (thus, HS^hi^ protein receptor^low^), and K562-VLDLR cells (**Supp. Fig. 3A-D**). Notably, despite its expression, HS is not a dominant factor for productive infection of K562-VLDLR cells as they are still highly infectable following heparinase II digestion (**Supp. Fig. 1B**). Therefore, the K562-VLDLR cells functionally act as complimentary HS^low^ VLDLR^high^ cells.

Sequencing of supernatants from the K562-VLDLR cells at passage 5 and 10 revealed that all three viruses retained the original mutant residues (**Table 1**), and by passage 5 the 71-77 and 156-157 mutants each acquired an additional charge-reversing substitution mutation (71-77-E244K and 156-157-E206K), with residues physiochemically similar to documented passage-associated changes in arboviruses found to increase HS-binding^7^. In addition, the 206K residue was reported to allow EEEV strain PE-6 to use an additional E2 domain B shelf VLDLR-binding site^11-13^. By passage 10, both replicates for the 84-119 mutant also acquired the E206K mutation. Sequencing of BHK-21 cell supernatants at passages 5 and 10 revealed the 71-77 and 156-157 mutants had each acquired a mutation to a positively charged amino acid while, again, retaining the original mutant residues (71-77-E147K and 156-157-Q8K) (**Table 1**). No mutations or reversions were observed for the 84-119 mutant (**Table 1**). Notably, all acquired mutations except for E244K are within three amino acid residues of MXRA8, LDLRAD3, and/or VLDLR-binding residues (**Supp. Fig. 2**) ^11-13,29,30,33^.

**Table 1.** Mutations acquired after passaging chimeric HS-binding mutants. Mutations that are different from the WT sequence are bolded, and mutations acquired during passage are bolded and underlined.

| SINV/EEEV eGFP TaV | E2 amino acid position |  |  |  |  |  |  |  |  |  |  |
| --- | --- | --- | --- | --- | --- | --- | --- | --- | --- | --- | --- |
|  | 8 | 71 | 74 | 77 | 84 | 119 | 147 | 156 | 157 | 206 | 244 |
| WT | Q | K | K | K | R | R | E | K | R | E | E |
| 71-77 parent | Q | **A** | **A** | **A** | R | R | E | K | R | E | E |
| 71-77 BHK5.1 | Q | **A** | **A** | **A** | R | R | <u>**K**</u> | K | R | E | E |
| 71-77 BHK5.2 | Q | **A** | **A** | **A** | R | R | <u>**K**</u> | K | R | E | E |
| 71-77 BHK10.1 | Q | **A** | **A** | **A** | R | R | <u>**K**</u> | K | R | E | E |
| 71-77 BHK10.2 | Q | **A** | **A** | **A** | R | R | <u>**K**</u> | K | R | E | E |
| 71-77 K562-VLDLR5.1 | Q | **A** | **A** | **A** | R | R | E | K | R | E | <u>**K**</u> |
| 71-77 K562-VLDLR5.2 | Q | **A** | **A** | **A** | R | R | E | K | R | E | <u>**K**</u> |
| 71-77 K562-VLDLR10.1 | Q | **A** | **A** | **A** | R | R | E | K | R | E | <u>**K**</u> |
| 71-77 K562-VLDLR10.2 | Q | **A** | **A** | **A** | R | R | E | K | R | E | <u>**K**</u> |
| 84-119 parent | Q | K | K | K | **A** | **A** | E | K | R | E | E |
| 84-119 BHK5.1 | Q | K | K | K | **A** | **A** | E | K | R | E | E |
| 84-119 BHK5.2 | Q | K | K | K | **A** | **A** | E | K | R | E | E |
| 84-119 BHK10.1 | Q | K | K | K | **A** | **A** | E | K | R | E | E |
| 84-119 BHK10.2 | Q | K | K | K | **A** | **A** | E | K | R | E | E |
| 84-119 K562-VLDLR5.1 | Q | K | K | K | **A** | **A** | E | K | R | E | E |
| 84-119 K562-VLDLR5.2 | Q | K | K | K | **A** | **A** | E | K | R | E | E |
| 84-119 K562-VLDLR10.1 | Q | K | K | K | **A** | **A** | E | K | R | **<u>K</u>** | E |
| 84-119 K562-VLDLR10.2 | Q | K | K | K | **A** | **A** | E | K | R | **<u>K</u>** | E |
| 156-157 parent | Q | K | K | K | R | R | E | **A** | **A** | E | E |
| 156-157 BHK5.1 | **<u>K</u>** | K | K | K | R | R | E | **A** | **A** | E | E |
| 156-157 BHK5.2 | **<u>K</u>** | K | K | K | R | R | E | **A** | **A** | E | E |
| 156-157 BHK10.1 | **<u>K</u>** | K | K | K | R | R | E | **A** | **A** | E | E |
| 156-157 BHK10.2 | **<u>K</u>** | K | K | K | R | R | E | **A** | **A** | E | E |
| 156-157 K562-VLDLR5.1 | Q | K | K | K | R | R | E | **A** | **A** | **<u>K</u>** | E |
| 156-157 K562-VLDLR5.2 | Q | K | K | K | R | R | E | **A** | **A** | **<u>K</u>** | E |
| 156-157 K562-VLDLR10.1 | Q | K | K | K | R | R | E | **A** | **A** | **<u>K</u>** | E |
| 156-157 K562-VLDLR10.2 | Q | K | K | K | R | R | E | **A** | **A** | **<u>K</u>** | E |

To test whether these additional mutations partially or fully restored HS-binding and/or VLDLR receptor usage, we determined the infectivity of the chimeric passaging mutants on CHO-K1 *versus* GAG- or HS-deficient CHO-pgsA-745 or CHO-pgsD-677 cells and the increase of infectivity for cells that overexpress the protein receptors *versus* control cells (**Fig. 3A, C-D**). Since the 156-157 mutant has more pronounced effects on these phenotypes compared to the 84-119 mutant (**Figs. 1A-B and 2A-C**), we evaluated the impact that the addition of the E206K mutation using the 156-157/206K mutant only. Both mutants that arose from passaging on BHK-21 cells (71-77/147K and 156-157/8K) exhibited significantly reduced infectivity for both CHO-pgsA-745 and CHO-pgsD-677 cells compared to CHO-K1 (**Fig. 3A**). Additionally, both mutants that arose from passaging on K562-VLDLR cells (71-77/224K and 156-157/206K), also exhibited significantly reduced infectivity for the GAG- and HS-deficient cells compared to the CHO-K1 controls (**Fig. 3A**). Therefore, selection on both HS^hi^ protein receptor^low^ and HS^low^ VLDLR^high^ cells yields mutations that increase HS-dependent infectivity, presuming that protein receptor-mediated infectivity is limited on CHOs and equal between mutant and WT cells. Both 71-77/147K and 156-157/8K mutants showed increased binding to CHO-K1 cells compared to their parent mutation (**Fig. 3B**). However, the GAG-dependent infectivity result was only recapitulated for cell binding with the chimeric 71-77/147K mutant, which exhibited a significant decrease in binding to CHO-psgA-745 cells compared to CHO-K1 (**Fig. 3B**). The 71-77/244K, 156-157/8K, and 156-157/206K mutants all exhibited no decrease in binding to CHO-psgA-745 *versus* CHO-K1 cells despite a significant decrease in infectivity (**Fig. 3A-B**). However, since the decreased infectivity was less pronounced for the 71-77/244K, 156-157/8K, and 156-157/206K mutants compared to the 71-77/147K mutant, it is possible that the binding assay may not be sensitive enough to detect subtle differences in binding that affect infectivity; or these mutations may allow for binding to a different structure on those cells that does not promote productive infection.

**Fig. 3.**
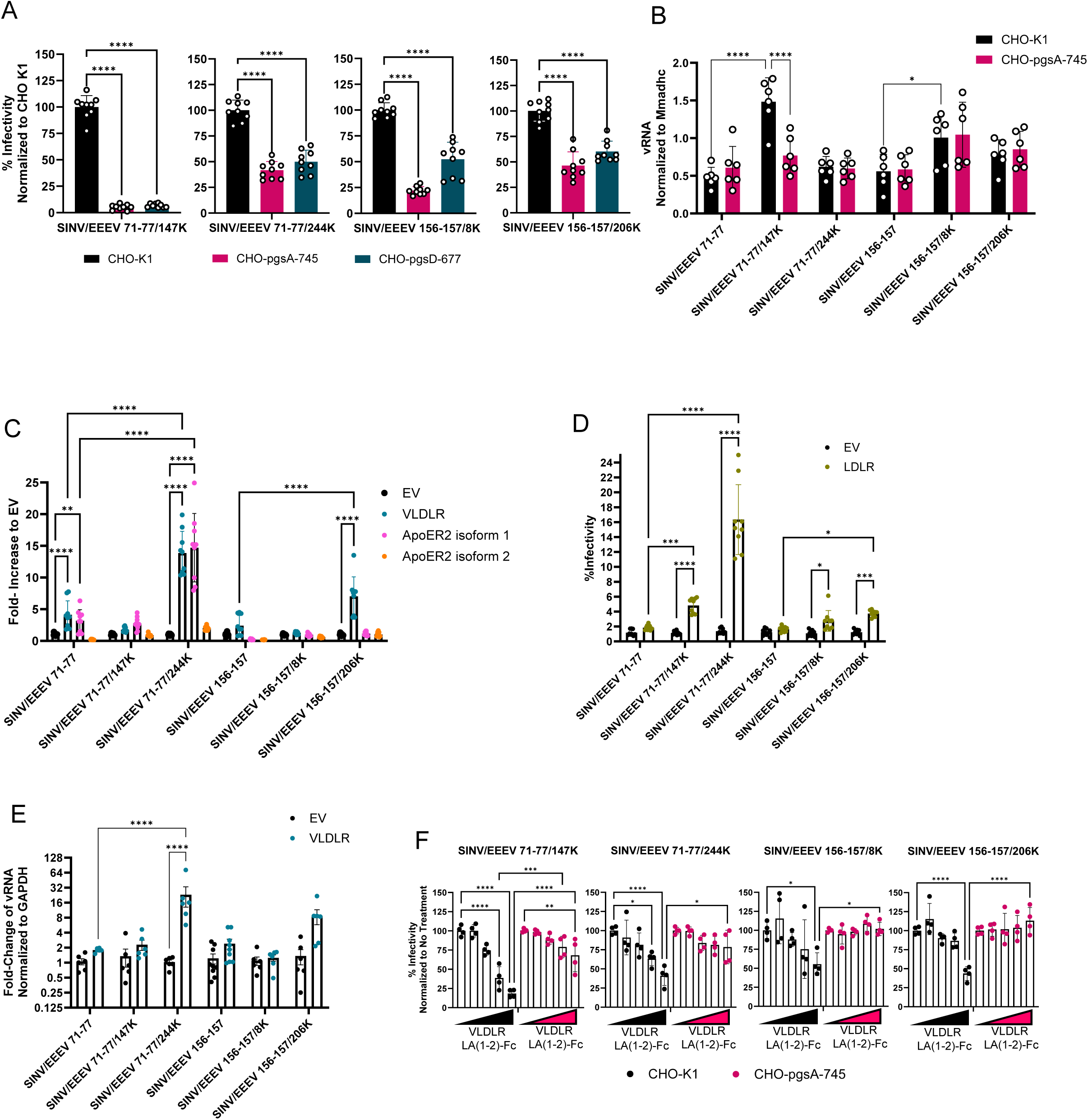
Passage of HS/protein receptor binding site mutants on cultured cells selects for mutations that impact binding of both receptors. **A** Infectivity of double SINV/EEEV mutants found in passaging experiment on CHO-K1, CHO-pgsA-745, or CHO-pgsD-677 cells. Relative infectivity compared to CHO-K1 was determined for all viruses (n = 9 from 3 independent experiments). Viruses were titrated on all three cell types, and percent infectivity compared to CHO-K1 was determined for each cell type. **B** Binding of chimeric passaging mutant viruses to CHO-K1 and CHO-pgsA-745 cells, performed as described in **Fig. 1B** (n = 6 from 2 independent experiments) with bound virus expressed as the ratio to *Mmadhc*. (Data for single mutant parent viruses are the same as in **Fig. 1B**.) **C-D** Infectivity of passaging mutants in (**C**) K562 cells expressing EV, VLDLR, ApoER2 isoform 1, or ApoER2 isoform 2 (n = 9 from 3 independent experiments, except for the 71-77 and 156-157 mutants on K562-EV cells where n = 15 from 5 independent experiments) or (**D**) THP-1 EV or LDLR cells (n = 9 from 3 independent experiments) quantified by flow cytometry. (Data for single mutant parent viruses are the same as in **Fig. 2A-B**.) **E** Binding of chimeric WT and mutant viruses to K562-EV or K562-VLDLR cells, performed as described in **Fig. 2C** (n = 6 from 2 independent experiments, except for 1561-57 where n= 9 from 3 independent experiments). Binding is expressed as a ratio to *GAPDH* and normalized to EV. (Data for single mutant parent viruses are the same as in **Fig. 2C**.) **F** Neutralization of chimeric passaging mutant viruses by VLDLR LA(1-2)-Fc (100-0.1µg in 10-fold dilutions and 0 µg control) in CHO-K1 and CHO-pgsA-745 cells (n = 4 from 2 independent experiments). Significance was determined by (**A**) one-way ANOVA with Bonferroni’s post-hoc tests and (**B-F**) two-way ANOVA with Tukey’s post-hoc tests. Error bars show SD, except for (**E**) where error bars show SEM; *(p<0.05), **(p<0.01), ***(p<0.001), ****(p<0.0001).

When protein receptor-dependent infection was assessed, of the mutants that arose during BHK-21 (HS^hi^ protein receptor^low^) cell passage, the 71-77/147K mutant exhibited increased infectivity *versus* the 71-77 parent only on THP-1 LDLR cells. Although exhibiting a small but significant infectivity increase for THP-1 LDLR cells *versus* THP-1 EV cells, the 156-157/8K mutant showed no significant difference in infectivity *versus* the 156-157 parent for any of theover-expression cells (**Fig. 3C-D**). For the mutants that arose during K562-VLDLR (HS^low^ VLDLR^high^) cell passage, the 156-157/206K mutant only showed increased infectivity for the K562-VLDLR and THP-1 LDLR cells compared to the 156-157 parent, whereas the 71-77/244K mutant showed increased infectivity compared to the 71-77 parent on K562-VLDLR, K562-ApoER2 isoform 1 and THP-1 LDLR cells (**Fig. 3C-D**). The infectivity results were recapitulated when virus binding to K562-EV and K562-VLDLR was assessed. The 71-77/147K and 156-157/8K mutants exhibited similar binding to their parental viruses with K562-EV and K562-VLDLR cells, while the 71-77/244K and 156-157/206K mutants exhibited increased binding to K562-VLDLR cells, although the increase was not significant (p=0.09) for the 156-157/206K mutant (**Fig. 3E**). These data suggest that mutations selected in the context of protein receptor abundance increase both HS- and protein receptor-dependent infection, whereas selection for mutations in the context of HS abundance leads to increased HS-binding and HS-dependent infection, and can, but does not always, lead to increased protein receptor utilization.

To determine whether increased HS-dependent infectivity/protein receptor interaction would also lead to increased protein receptor decoy neutralization, we tested the ability of VLDLR LA(1-2)-Fc to neutralize the passaging mutants on CHO-K1 and CHO-pgsA-745 cells. Both of the 71-77 passaging mutants showed comparable, limited neutralization as observed for the 71-77 parent on CHO-pgsA-745 cells. However, unlike the parental 71-77 virus, both mutants were significantly neutralized by higher concentrations of VLDLR LA(1-2)-Fc when GAGs were present (**Figs. 2E and 3F**). Similarly, both 156-157 passaging mutants showed no neutralization on GAG-deficient cells but were significantly neutralized at the 100 µg/mL concentration on CHO-K1 cells (**Fig. 3F**). These results suggest that in the context of low-to-negative protein receptor expression, the decoy can block infectivity through the blocking of HS interactions. However, the role of the limited and equal VLDLR receptor expression on CHO-K1 or CHO-pgsA-745 cells (**Supp. Fig. 3B**) is not clear in these experiments. Together, mutant data indicate that single-site, positively charged amino acid mutations selected during passage can simultaneously increase interactions with HS and protein receptors.

### Mutation of HS/protein receptor binding sites attenuates EEEV disease and diminishes protection conferred by receptor decoy inhibitors

To determine whether the HS-binding site mutations led to attenuation following the natural route of vertebrate infection, mice were infected with WT or mutant EEEV strain FL93-939 viruses with genomes equivalent to 10^3^ WT PFU subcutaneously (sc.) in the rear footpad (fp.). As with WT, infection with the 84-119 mutant led to rapid mortality and limited clinical signs (**Fig. 4A-C**). As previously reported^3,4^, the 71-77 mutant led to earlier signs of clinical illness and a significantly extended average survival time (AST) compared to the WT. Despite the intermediate *in vitro* phenotype, the 156-157 mutant was the most attenuated, resulting in a significantly longer AST compared to the WT and the other mutants (p<0.0001 compared to WT, 71-77, and 84-119, log-rank test), but it was still uniformly lethal (**Fig. 4A**).

**Fig. 4.**
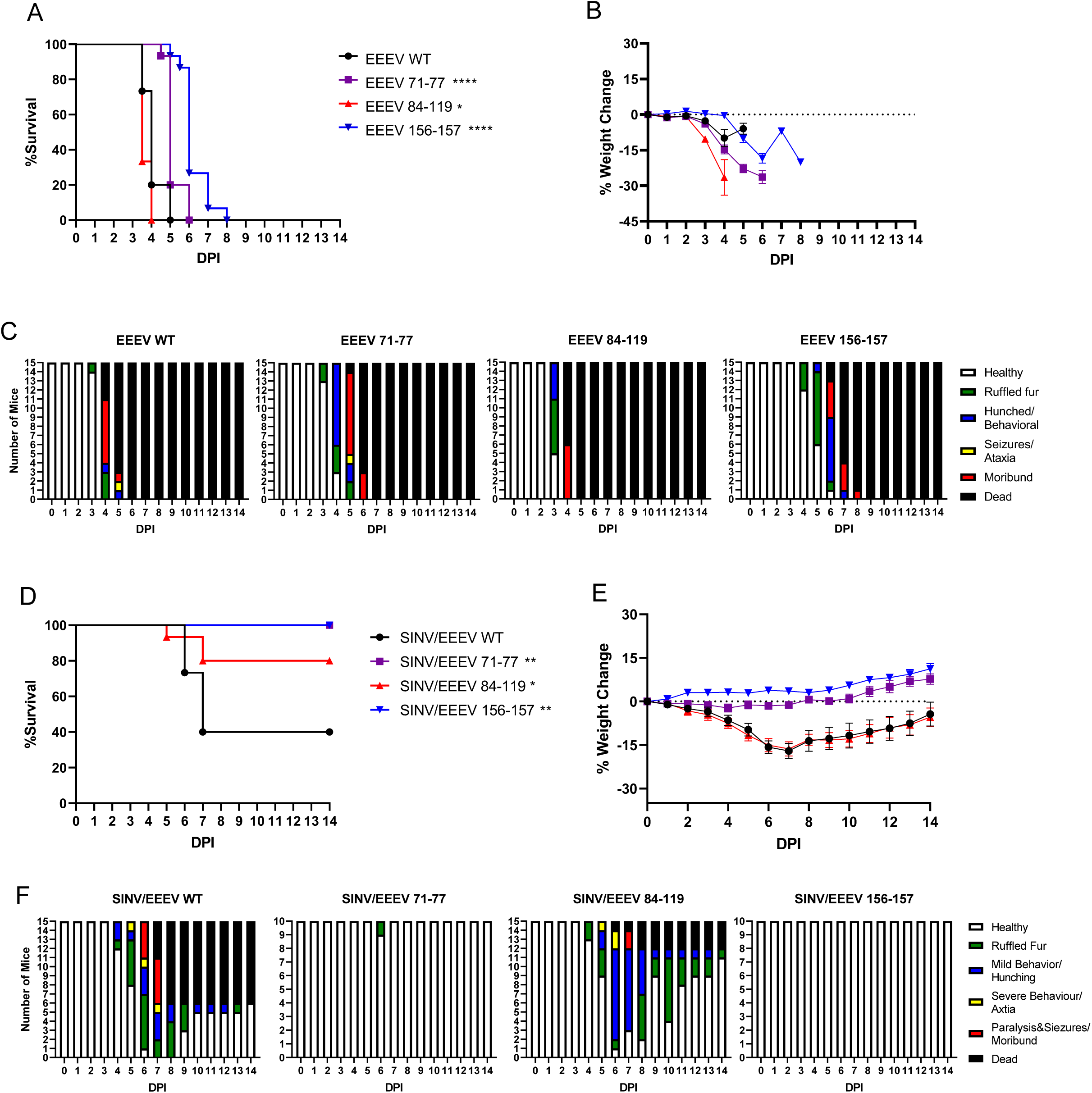
Mutation of HS/protein receptor binding sites attenuates EEEV disease. **A-C** Groups of five CD-1 mice were infected with equal genomes of WT and mutant EEEV viruses equivalent to 10^3^ WT PFU in fp. (n = 15 mice from 3 independent experiments). **A** Survival. **B** Weight change. **C** Clinical signs. **D-E** Groups of five CD-1 mice were infected ic. with equal genomes of WT and mutant SINV/EEEV virus equivalent to 10^4^ WT PFU (n = 15 mice from 3 independent experiments for SINV/EEEV WT and SINV/EEEV 84-119, and n= 10 mice from 2 independent experiments for SINV/EEEV 71-77 and SINV/EEEV 156-157). **D** Survival. **E** Weight change. **F** Clinical signs. All error bars show SEM. Significance determined by log-rank test; *(p<0.05), ** (p<0.01) ***(p<0.001), ****(p<0.0001).

To provide a more sensitive measurement of neurovirulence in the absence of replication outside the CNS, we infected mice intracranially (ic.) with equal genomes of SINV/EEEV chimeric viruses equivalent to 10^4^ PFU of the WT. The chimeras are attenuated in mice *versus* parental EEEV viruses^34^. Mice infected with the 71-77 or 156-157 mutants did not develop lethal encephalitis and showed no or limited clinical signs of disease (**Fig. 4D-F**). Although 80% of mice infected with the 84-119 mutant survived, all developed clinical signs of disease and had similar weight loss as mice infected with the WT (**Fig. 4E-F**) and the 84-119 mutant resulted in only 20% mortality, which was significantly lower than the 60% mortality rate of the WT chimera (**Fig. 4D**).

Recently we, with collaborators, demonstrated that pre-treatment with VLDLR LA(1-2)-Fc protects mice from lethal sc. infection of WT EEEV^11^. Due to the significant decreases in infection on cells over-expressing protein receptors and the significant differences noted in the ability of VLDLR LA(1-2)-Fc to neutralize the mutant viruses, we tested the ability of VLDLR LA(1-2)-Fc to protect mice from infection with genome equivalents of EEEV WT and the E2 mutants. Mice were treated intraperitoneally with 100 µg of either VLDLR LA(1-2)-Fc or LDLRAD3 LA1-Fc control six hours before infection. All mice treated with the LDLRAD3 LA1-Fc control before infection succumbed to infection (**Fig. 5**). As with previous studies^11^, mice pretreated with VLDLR LA(1-2)-Fc were almost completely protected from WT infection (**Fig. 5A and Supp. Fig. 4**). Despite the significant reduction *versus* WT in VLDLR-mediated infectivity and neutralization *in vitro* (**Fig. 2A, D-E**), mice were also largely protected from the 71-77 mutant following pretreatment with VLDLR LA(1-2)-Fc (**Fig. 5B**), suggesting that the decoy does bind to the 71-77 mutant virus. Pretreatment with VLDLR LA(1-2)-Fc led to partial protection from lethal disease with an extended AST compared to controls for the 84-119 mutant (**Fig. 5C**). In line with the *in vitro* data showing no binding, the 156-157 mutant showed no difference in outcome with pretreatment of LDLRAD3 LA1-Fc or VLDLR LA(1-2)-Fc (**Fig. 5D**).

**Fig. 5.**
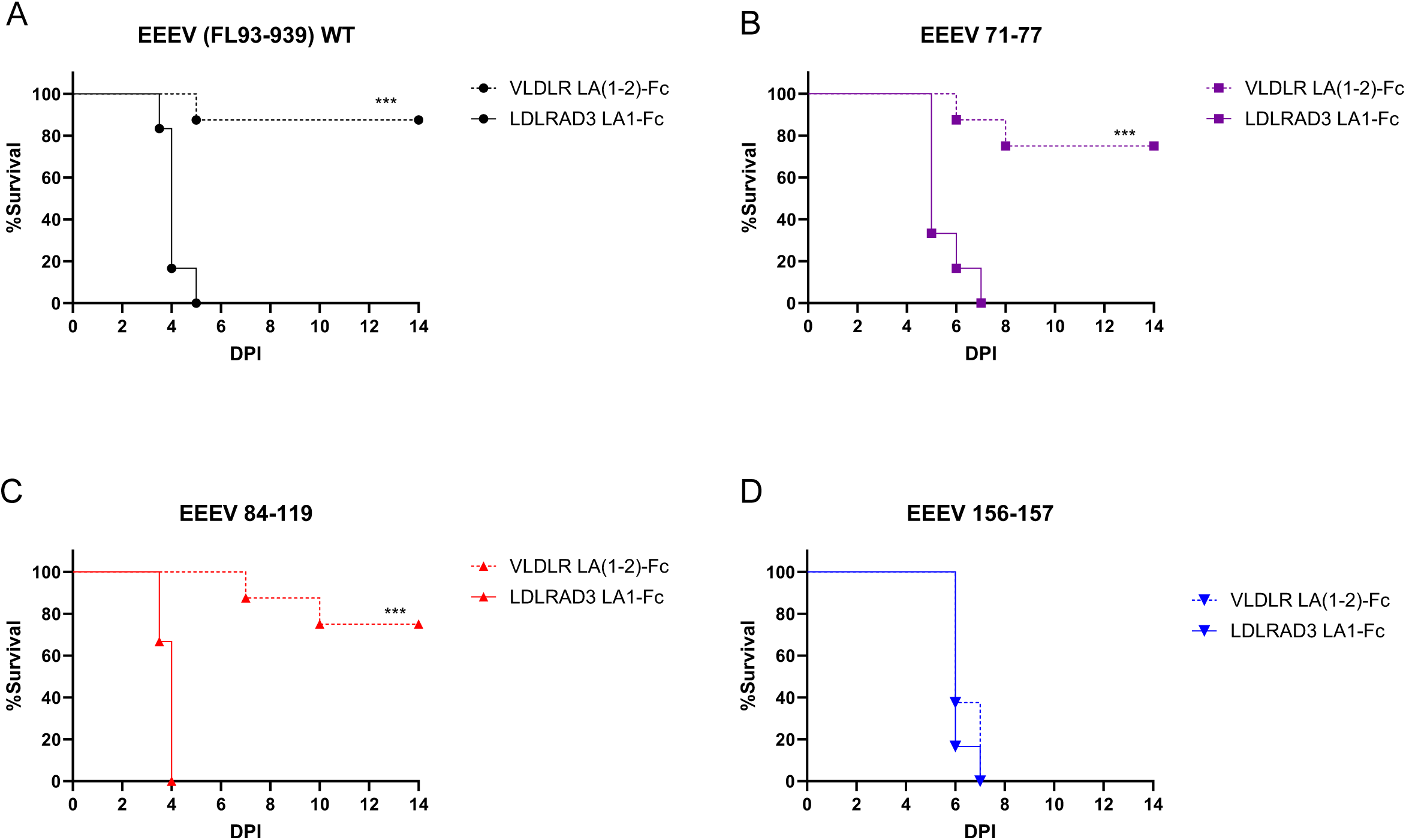
Mutation of HS/protein receptor binding sites diminishes protection conferred by the receptor decoy inhibitor. Survival data for mice that were treated intraperitoneally with 100 µg of VLDLR LA(1-2)-Fc (dashed line) or LDLRAD3 LA1-Fc (solid line) as a control. Six hours after treatment, mice were infected fp. with equal genomes of WT and mutant EEEV viruses equivalent to 10^3^ WT PFU (n = 6 mice from 2 independent experiments for LDLRAD3 LA1-Fc and n = 8 mice from 2 independent experiments for VLDLR LA(1-2)-Fc). **A** EEEV WT (FL93-939). **B** EEEV 71-77. **C** EEEV 84-119. **D** EEEV 156-157. Significance of survival for mice treated with control *versus* VLDLR LA(1-2)-Fc was determined by log-rank test; *(p<0.05), **(p<0.01), ***(p<0.001).

### Mutation of HS/protein receptor binding sites diminishes transmission to and dissemination within mosquito vectors

Finally, we examined the effects of HS-binding site mutations on infection and dissemination within a mosquito vector. To determine whether the E2 mutants would lead to a restriction in productive infection, measured by the detection of the virus in the body, or dissemination, measured by the detection of the virus in the heads, we infected *Aedes albopictus* mosquitoes, an EEEV bridge vector species^35,36^ with equal genomes of WT or HS-binding site mutants *via* artificial bloodmeals (**Table 2**). All mosquitoes infected with the WT had detectable virus in their bodies and heads. Infection with the 84-119 mutant showed a small decrease in infection and a modest but significant decrease in dissemination, with 90% showing infection in bodies and 80% showing infection in heads. In contrast, the 71-77 mutant exhibited a significant decrease in both infection and dissemination, with only 20% of mosquitoes having virus in the body, and 10% in their heads (**Table 2**). Strikingly, none of the mosquitoes infected with the 156-157 mutant were productively infected, suggesting that the E2 K156-R157 residues are critical for establishing infection in mosquitoes.

**Table 2.** Mutation of HS/protein receptor binding sites diminishes transmission to and dissemination within mosquito vectors. *Aedes albopictus* mosquitoes (n = 20 mosquitoes) were infected via artificial bloodmeal with equal genomes of WT or mutant EEEV equivalent to 8.2 log_10_ WT PFU/mL. At 7 dpi mosquitoes were collected, and the presence of virus in the heads and bodies was determined by infectious assay. Significance was determined by two-way ANOVA with Dunnett’s post-hoc tests.

|  | Body (Infection) |  | Head (Dissemination) |  |
| --- | --- | --- | --- | --- |
|  | # infected/total (%) | p-value | # infected/total (%) | p-value |
| EEEV WT | 20/20 (100%) | - | 20/20 (100%) | - |
| EEEV 71-77 | 4/20 (20%) | <0.0001 | 2/20 (10%) | <0.0001 |
| EEEV 84-119 | 18/20 (90%) | 0.4663 | 16/20 (80%) | 0.0395 |
| EEEV 156-157 | 0/20 (0%) | <0.0001 | 0/20 (0%) | <0.0001 |

## DISCUSSION

To date, EEEV remains the only alphavirus for which unpassaged strains have been shown to bind to HS efficiently^3^. Previous work mapped positively charged residues at positions 71, 74, and 77 of the E2 glycoprotein of EEEV, as critical for EEEV HS-binding and contribute to neurovirulence and reduction of draining lymph node infection and viral prodromal disease^3,4^. Studies using cryo-EM and 6kD heparin further identified heparin/HS-binding sites coordinated by positively charged residues at E2 84-119 and 156-157^8^. Our current studies demonstrate that the newly identified E2 residues also contribute to EEEV-HS interactions that promote infection *in vitro* and affect pathogenesis *in vivo* (**Table 3**). Mutation of the 71-77 site, as previously reported^3,4^, resulted in a virus that was significantly reduced *versus* the WT in all assays of HS-dependence. Mutation of the 156-157 site altered infectivity dependence with increasing ionic strength, yielded variable dependence on cellular HS expression for infectivity measured with GAG/HS-deficient cells and heparinase digestion, and decreased binding to heparin. The 84-119 site appears to alter EEEV-HS interactions but was significantly different from WT in only ionic strength assays. Therefore, while differing in quantitative and qualitative aspects of the HS interaction, each site contributes to the HS-dependence of EEEV infection.

**Table 3.**
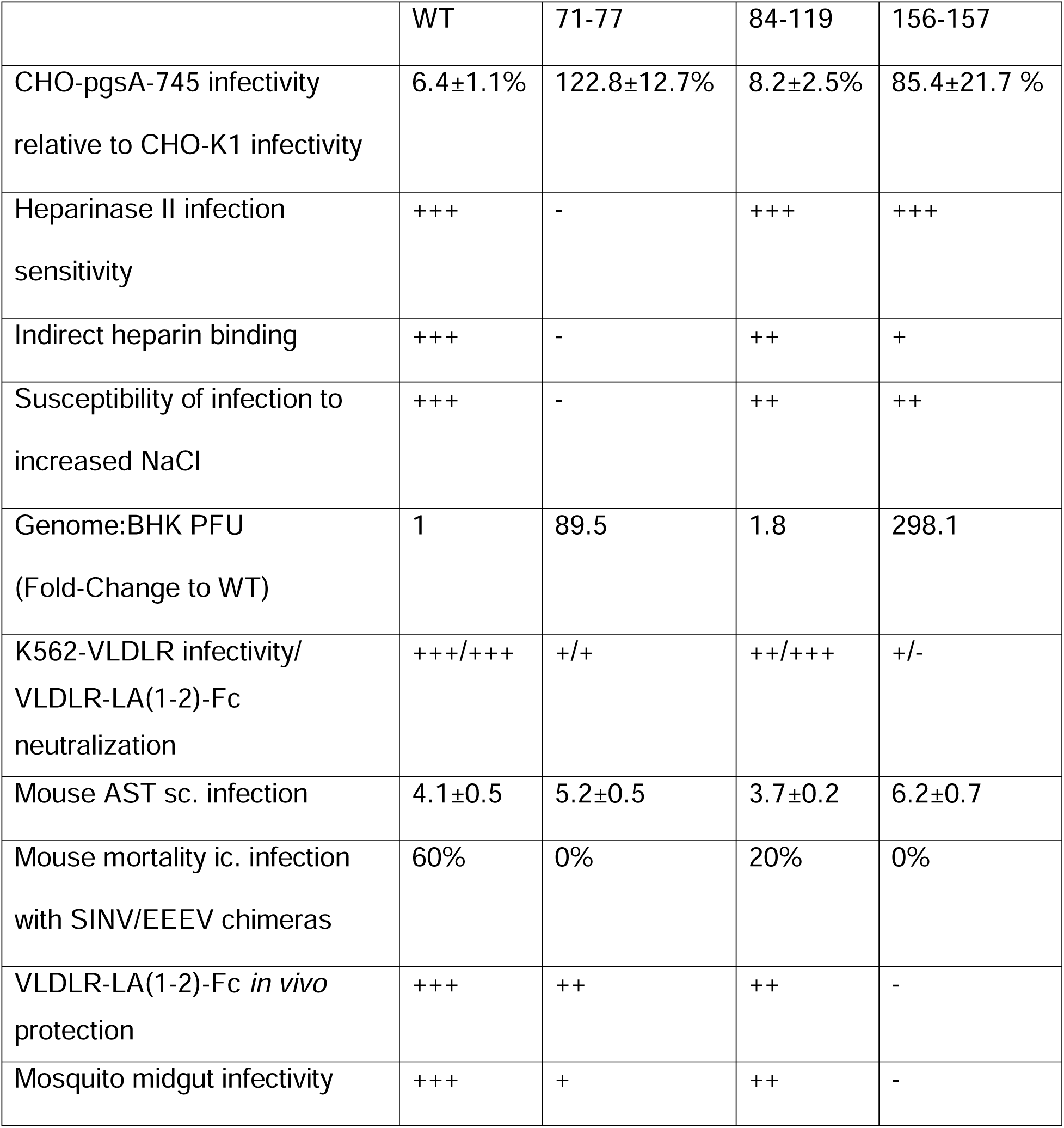
Summary of effects of each mutation on EEEV HS or receptor interactions. Plus and minus signs represent a subjective ranking of the relative magnitude of each phenotype (- = no activity, + - +++ represents a continuum from low to high magnitude). AST = average survival time; sc. = subcutaneous; ic. = intracranial.

Recently, the residues that mediate EEEV-VLDLR binding were mapped in cryo-EM binding studies. Two of the E2 residues that were identified as being critical for engagement with the VLDLR LA1 domain were K156 and R157^11-13^. The K156 residue was also shown to be important for EEEV-ApoER2 isoform 2 interactions^12^. Consistent with these new reports^11-13^, our data indicates that charged-to-alanine mutation at the 156-157 HS-binding site ablates EEEV-VLDLR interactions and ApoER2- and LDLR-dependent infectivity. This finding is notable as it shows that two mutations are sufficient to ablate EEEV interaction with multiple receptors, which would need to be considered for the development of infection-blocking reagents as therapeutics. Despite not being identified as VLDLR contact residues^11-13^, the 71-77 mutation also significantly impacted EEEV interactions with VLDLR, ApoER2, and LDLR, albeit less than the 156-157 mutation. It is possible that the 71-77 mutation impacts the ability of EEEV to engage with the LA2 domain of VLDLR, as the residues that participate in the domain A ‘shelf’ engagement with LA2 are in proximity to the 71, 74, and 77 residues (**Fig. 6**)^11-13^. The 84-119 mutant, whose altered residues were also not identified as VLDLR contacts^11-13^, had a minor impact on VLDLR- and ApoER2 isoform 1-dependent infectivity and a significant reduction of ApoER2 isoform 2- and LDLR-dependent infectivity, implying a more limited and specific effect on protein receptor interactions.

**Fig. 6.**
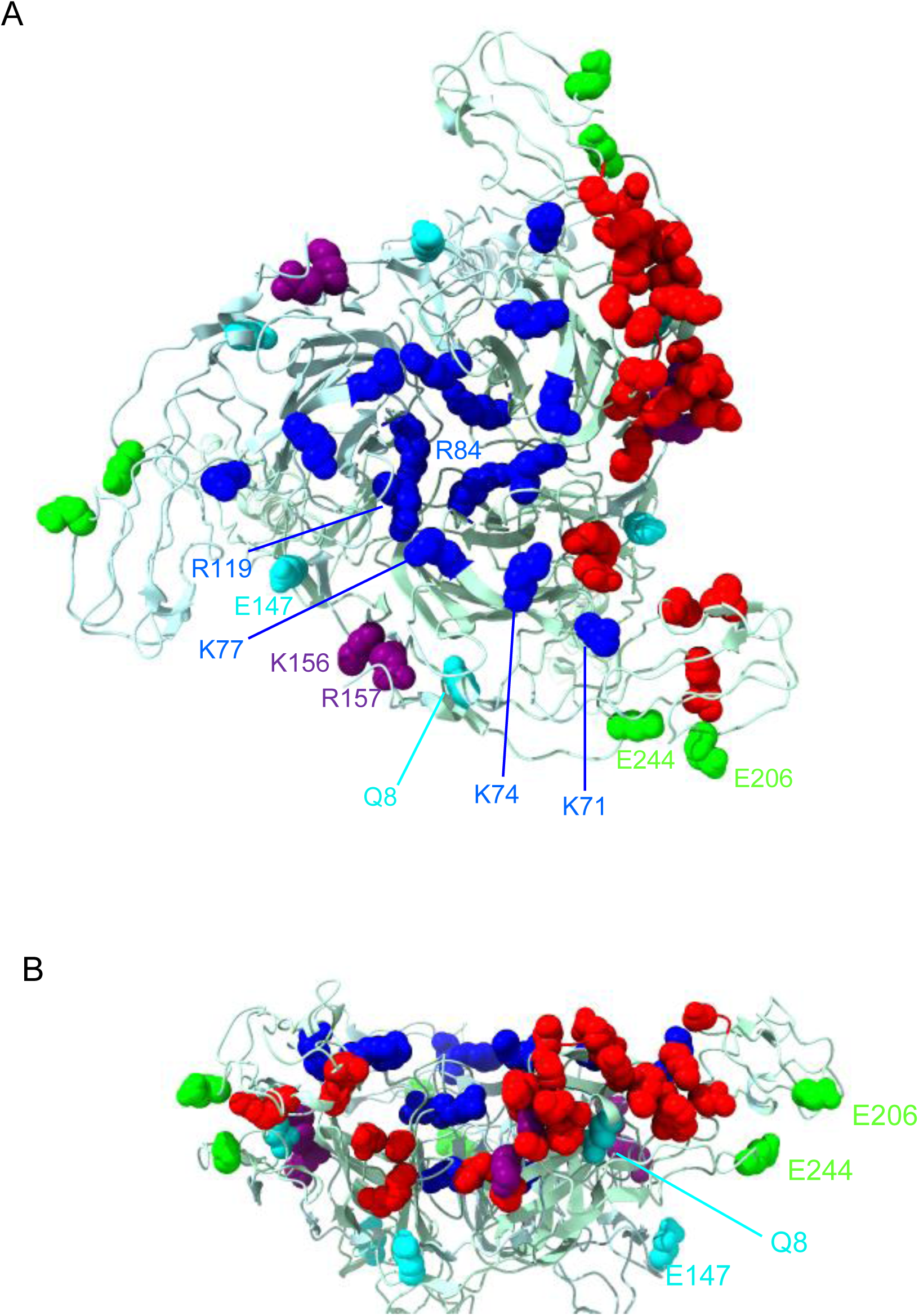
Location of protein receptor- and HS-binding residues on a ribbon model of the EEEV E2 trimer. A ribbon model of the structure of EEEV E2 proteins as they appear in the E1/E2 trimeric spike. Side chains are displayed for residues involved in HS-binding (blue), residues identified in structural analysis as being involved in binding to one VLDLR molecule^11-13^ (red), and residues that are involved in both HS-binding and are direct contacts for VLDLR-binding (purple). Side chains for residues mutated during *in vitro* passage are lime green (K562-VLDLR) or cyan (BHK-21). **A** Top view. **B** Side view. Figures were made using UCSF ChimeraX^43,44^.

A summary of the relative effects of the HS binding site mutations (71-77, 84-119, and 156-157) on HS and protein receptor utilization (**Table 3**) suggests that EEEV HS- and protein receptor-binding overlap with each of the three E2 sites. The 84-119 site has a modest effect on both phenotypes, whereas the 71-77 and 156-157 sites have more profound, although phenotypically different, effects. The 156-157 site has direct contact with VLDLR ^11-13^, and the 156-157 mutant appears to have little to no interaction with any identified protein receptor. Further evidence for the overlap of HS- and protein receptor-binding by EEEV is provided in the phenotypes of cell passaging mutants. When EEEV was passaged on HS^high^ protein receptor^low^ cells, both acquired mutations (E2 8K and 147K) increased EEEV-HS interactions, with the 147K mutation also increasing LDLR-dependent infectivity. However, when passaged on cells that have a HS^low^ VLDLR^high^ phenotype, the adaptive E2 206K and 244K mutations (located in domain B, which engages VLDLR)^11-13^ increased both protein receptor and HS interactions. Furthermore, our data support the observation that lysine at E2 206 results in a new site of VLDLR engagement (as found with the PE-6 strain of EEEV)^11-13^ when the lysine at E2 156 is mutated, and may suggest that this is a cell-adaptive change.

In further support of a link between EEEV engagement with HS and protein receptors, neutralization of chimeric WT virus and passaging mutants by the VLDLR LA(1-2)-Fc receptor decoy was greater on cells that expressed HS than on GAG-deficient cells. This suggests that the strong neutralizing phenotype of the VLDLR decoy is likely also due to inhibiting EEEV-HS interactions. In reciprocal experiments using heparin to neutralize EEEV infectivity, we observed significant competition of infection of VLDLR, ApoER2, or LDLR over-expressing cells, even when cells were genetically altered to lack HS. Since cryo-EM studies show that related and unrelated viruses have converged to utilize the same structural approach to engage with members of the LDL-receptor family^11,12^, it is possible that protein receptor engagement and HS-binding sites may overlap structurally and chemically. This may explain the seemingly unlikely phenomenon of single positively charged amino acid mutations conferring efficient HS binding to cell passaged viruses^7^, with these mutations often appearing adjacent to or to directly overlapping identified protein receptor-binding residues (**Supp. Fig. 2**). Indeed, there may be a balance during virus transmission such that satisfactory binding with protein receptors is achieved with minimal HS interactions but upon passage *in vitro*, where protein receptor abundance may be variable, single site mutations can rapidly confer increased cell attachment, thus rescuing infectivity. *In vivo*, selective pressures likely drive reductions in HS-binding due to enhanced clearance associated with the phenotype^18,37,38^. However, effects on protein receptor engagement are less predictable. It is notable that the 71-77 region is a hotspot for variation among EEEV strains, and such variants have altered engagement of HS receptors *versus* EEEV FL93-939^4^. Testing of VLDLR utilization by such variants is in progress.

Despite significantly decreasing EEEV-HS interactions and ablating EEEV interactions with all four identified protein receptors and mosquito infectivity, the 156-157 mutant was still uniformly fatal in mice. This result implies that loss of HS- and VLDLR/ApoER2/LDLR-binding via single-site mutation is not highly attenuating to EEEV in mammals, and it predicts the existence of additional attachment/entry receptors that have yet to be identified. Yet VLDLR LA(1-2)-Fc protects mice from lethal WT EEEV disease after sc. infection^11^, suggesting an important role for the alphavirus VLDLR (and other protein receptor) binding domains *in vivo*.

The abrogation of HS-binding diminished the neurovirulence of each mutant and enhanced neurovirulence has been attributed to the HS-binding phenotype^3^. Our observations that the mutations to EEEV also impact the utilization of protein receptors bring into question the role of each type of receptor in each of these phenomena. The decreased murine neurovirulence associated with diminution of HS- and VLDLR-interactions will require additional investigation to parse the different receptor engagement effects. The role of HS *versus* protein receptor binding upon suppression of lymphoid tissue infection and viremia, previously attributed to HS binding alone^3,7^, will also need to be determined. The role of VLDLR/ApoER2/LDLR may be clearer with mosquito infection, as complete loss of infectivity mediated by these receptors with the 156-157 mutant abrogates blood meal infection. Thus, positive charges at 156-157 are required for mosquito receptor engagement suggesting a receptor binding modality similar to that described for the identified protein receptors.

Since EEEV has evolved to circulate in nature with efficient HS-binding, it is unclear how applicable the overlap between HS- and protein receptor-binding may be to weakly HS-binding viruses or viruses that engage receptors outside of the LDL-receptor family. Testing of the interactions of WT and cell-culture adapted, HS-binding viruses (e.g., derived from VEEV^18^ or CHIKV^22^) with their cognate receptors LDLRAD3^19,30,31^ and MXRA8^29,32,33^ will be required to clarify this issue. Since LDL binding proteins have been identified as receptors for a number of arboviruses^39-42^ and *in vitro* acquisition of HS-binding appears to be a ubiquitous phenotype in this group^7^, the interplay between HS- and protein receptor-binding may be an important feature of the arbovirus-cell interaction.

## Acknowledgements

We thank Colleen Petersen, Shauna Vasilatos, Claudia Brelsford, and Ruimei Yun for their technical assistance. We acknowledge Jonathan Abraham from Harvard Medical School for a gift of the K562 cells expressing identified protein EEEV receptors. These studies were supported by NIH Grants R01 AI095436 and R01 AI153209 awarded to W.B.K., and R01 AI164653 to M.S.D., DTRA contract MCDC 2103-011 to W.B.K and M.S.D., M.D.H.A was supported by NIH/NIAID T32 AI049820.

## Author Contributions

M.D.H.A, S.C.W., and W.B.K designed the study. M.D.H.A., T.C.G., and C.C. performed experiments. M.S.D. and S.R. developed transgenic cells and produced the decoy receptors. C.S. designed and generated viral constructs and primers. T.L. designed and optimized primers. S.C.W. oversaw mosquito studies. W.B.K. obtained funding. M.D.H.A and W.B.K wrote initial draft, all other authors provided comments.

## METHODS

### Cells and viruses

Baby hamster kidney (BHK-21 [ATCC CCL-10]) cells were maintained in RPMI-1640 and supplemented with 10% heat-inactivated donor bovine serum (DBS; Gibco) and 10% tryptose phosphate broth (Moltox). Chinese hamster ovary (CHO)-K1 [ATCC CCL-61], GAG-deficient mutant CHO-pgsA-745 [ATCC CRL-2242], and HS-deficient mutant CHO-pgsD-677 [ATCC CRL-2244] cells were maintained in Ham’s F-12 medium supplemented with 10% heat-inactivated fetal bovine serum (FBS, Gibco). Human lymphoblast K562 cells expressing VLDLR, ApoER2 isoform1, ApoER2 isoform2, or empty vector (EV) controls^9^ were kindly provided by Jonathan Abraham (Harvard Medical School, Boston) and maintained in RPMI-1640 with 10% heat-inactivated FBS, 20 mM HEPES (Corning), and 2 µg/mL puromycin (Mirus). African green monkey Vero [ATCC CCL-81] and N2a ΔB4galt7ΔLDLR-TC LDLR (N2a ΔB4galt7-LDLR) cells^10^ were maintained in Dulbecco’s modified Eagle’s medium (DMEM) supplemented with 10% heat-inactivated FBS (Omega Scientific), the N2as also being supplemented with 10 mM HEPES. THP-1 cells expressing LDLR or EV controls^10^ were maintained in RPMI-1640 with 10% heat-inactivated FBS (Omega Scientific), 10 mM HEPES, and 200 µg/mL hygromycin (Fisher). All media contained 2mM L-glutamine, 100 U/mL penicillin, and 0.5 mg/mL streptomycin.

The following viruses were used: SINV/EEEV (FL93-939), SINV/EEEV E2 71-77, SINV/EEEV E2 84-119, SINV/EEEV E2 156-157, SINV/EEEV E2 71-77/147K, SINV/EEEV E2 71-77/244K, SINV/EEEV E2 156-157/8K, SINV/EEEV E2 156/157/206K, EEEV (FL93-939), EEEV E2 71-77, EEEV E2 84-119, and EEEV E2 156-157. EEEV strain FL93-939 was generated from a cDNA clone^45^. E2 mutants EEEV 71-77^3^, EEEV 84-119^8^, and EEEV 156-157^8^ were constructed as previously described using the QuikChange II XL mutagenesis kit (Agilent Technologies). Fragment-swapping strategies were also used to generate combinations of the E2 mutations by the corresponding restriction sites (Mlu I, EcoR I, and Not I); constructs were confirmed by Sanger sequencing carried out by Genewiz. Chimeric mutant viruses were generated by replacing the structural proteins of SINV strain TR339 with the indicated virus as previously described^20,46^. eGFP and Nano-luciferase (nLuc) TaV reporter viruses were constructed as described in Sun *et al*^46^.

### Susceptibility to absence of GAGs and high ionic conditions

Viruses were serially diluted in virus diluent (VD; PBS containing cations supplemented with 1% DBS, 100 U/mL penicillin, and 0.5 mg/mL streptomycin) and titrated in triplicate on control CHO-K1, CHO-pgsA-745, and CHO-pgsD-677 cells for 1 h at 37 C. After infection, cells were overlayed with immunodiffusion-grade agarose (MP Biomedical) and incubated at 37°C 5% CO_2_ for 18-20 hpi, and then GFP-expressing plaques were counted by fluorescence microscopy. Titers were determined on all cell types, and percent infectivity compared to the CHO-K1 controls was determined for each virus. To test whether mutant viruses are susceptible to increasing ionic conditions, viruses were serially diluted in RPMI-1640 with increasing concentrations of NaCl. Diluted viruses were then used to infect BHK-21 cells for 1 h at 37 C in triplicate before being overlayed with immunodiffusion-grade agarose and incubated at 37°C 5%CO_2_ for two days. GFP-expressing plaques were counted by fluorescence microscopy and percent infectivity compared to RPMI-1640 only control, which has a salt concentration of approximately 102 mM, for each virus.

### Heparinase sensitivity assays

Heparinase treatments for BHK-21 cells were done as previously described^4,22^. Cells were washed once with PBS containing cations, then treated with 0, 0.5, or 1 U/mL heparinase I, II, or III (New England BioLabs) in PBS for 1.5h at 37 C. Cells were washed twice with PBS, infected with chimeric eGFP-expressing virus in triplicate for 1 h at 37 C, and then overlayed with immunodiffusion-grade agarose. At 18-20 hpi, GFP-expressing plaques were visualized by fluorescence microscopy. Titers for each treatment were determined, and percent infectivity compared to no treatment was determined for each condition.

For K562-EV and K562-VLDLR, 10^5^ cells were washed with PBS and then treated with 0, 1, 2, or 4 U/mL heparinase II in PBS for 1 h at 37 C. Cells were washed twice with PBS, infected with chimeric eGFP-expressing WT virus in triplicate for 1 h at 37 C, and then overlayed with media. At 20 hpi, cells were washed and fixed in 4% PFA and analyzed using BD Fortessa and FlowJo software. Percent infectivity was determined for every condition on both cell types.

### Indirect heparin binding assay

Viruses were serially diluted to approximately 10^6^ PFU/mL, then 50 µL of virus was added in triplicate to collagen- or heparin-agarose (Sigma) which had been washed with RPMI-1640 and incubated on ice for 30 minutes. Beads were pelleted, and the unbound virus in the supernatant was titrated in duplicate on BHK-21. Virus was allowed to infect the cells for 1 h at 37 C, and then cells were overlayed with immunodiffusion-grade agarose. At 18-20 hpi, GFP-expressing plaques were visualized by fluorescence microscopy. Titers for each virus in each condition were recorded, and the percent infectivity of the virus incubated with heparin beads compared to collagen beads was determined.

### Cellular binding assays

For CHO-K1 and CHO-pgsA-745, cells were seeded into a 96-well plate the previous day, and for K562-EV and K562-VLDLR cells 10^5^ cells were transferred to a v-bottom plate. Cells were chilled on ice and inoculated with 5×10^9^ genomes of the WT and mutant chimeric eGFP reporter viruses for 75 minutes on ice. Cell monolayers were then washed with chilled PBS before lysis by Tri-Reagent. RNA was isolated using 1-bromo-3-chloropropane (BCP) and isopropanol, and cDNA were generated using an M-MLV kit (Invitrogen). Binding to CHO cells was determined by digital droplet PCR (ddPCR) using the 2X ddPCR Supermix for Probes (No dUTP) kit (Bio-Rad) with primers targeting SINV nsp2 and *Mmadhc*^47^ (**Supp. Table 1**). Fold-change was directly quantified, and then percent binding compared to CHO-K1 was determined for each virus. Binding to K562 cells was determined by qPCR using TaqMan Fast Universal PCR Master Mix (2X), no AmpErase (Applied Biosystems) using primers targeting SINV nsp2 and *GAPDH* (**Supp. Table 1**).

### Replication curves

WT and mutant viruses were diluted in VD to genome equivalents corresponding to WT EEEV MOI 1 and used to infect BHK-21 cells in triplicate at 37°C for 1 hour. Then, media was added to the infected cells. At 0, 6, 12, 24, and 48 hpi, one-tenth of the media was collected, aliquoted, and stored at -80°C. The media on the cells was replenished with an equal volume of fresh media. Titers of collected samples were determined by plaque assay on BHK-21 cells.

### Infection of K562 and THP-1 cells

K562 cells expressing EV, VLDLR, ApoER2 isoform 1, or ApoER2 isoform 2 were infected with genome equivalents of WT or mutant SINV/EEEV viruses containing eGFP TaV at MOI 2.4 for WT. THP-1 cells expressing EV or LDLR were infected with 1:2 dilutions of stock eGFP TaV expressing SINV/EEEV WT or mutant viruses. Resuspended cells were spread across three wells and incubated at 37°C 5% CO_2_ for 1 h, then 100 µL media was added to wells. At 16-19 hpi, cells were washed and fixed in 4% PFA and analyzed using a BD Fortessa and FlowJo software. Percent infectivity was determined for each virus on each cell type.

### VLDLR decoy inhibition assays

Chimeric eGFP expressing WT or HS mutant viruses were diluted in VD and incubated with dilutions of VLDLR LA(1-2)-Fc or LDLRAD3-LA1-Fc as a control for 1 h at 37°C. Complexes were used to infect duplicate wells of Vero, CHO-K1, or CHO-pgsA-745 cells for 1 h at 37°C 5% CO_2_ before being overlayed with immunodiffusion-grade agarose. At 16-20 hpi, eGFP-expressing plaques were counted, and percent neutralization for each condition was determined for each virus.

### Heparin inhibition assays

Chimeric eGFP expressing WT or 71-77 mutant viruses were diluted 1:3 in VD and incubated with dilutions of Heparin or BSA as a control for 1 h at 37°C. Complexes were used to infect duplicate wells of N2a ΔB4galt7ΔLDLR-TC LDLR cells or 10^6^ cells of K562-VLDLR, K562-ApoER2, or THP-1 LDLR for 1h at 37°C 5% CO_2_ before being overlayed with immunodiffusion-grade agarose. At 16-20 hpi, N2A cells were detached with TrypLE, and all cells were washed and fixed in 4% PFA and analyzed using a BD LSR Fortessa and FlowJo software.

### *In vitro* virus passaging

Stocks of eGFP-expressing chimeric single mutant viruses were serially passaged ten times in duplicate on BHK-21 and K562-VLDLR cells. For BHK cells, stock or supernatant from previous passage was diluted 1:100 for passages 1 and 2, then 1:1,000 for passages 3-10. For K562-cells, stock or supernatant from the previous passage was diluted 1:100 or undiluted for all passages. Viral RNA from supernatant collected at passages 5 and 10 from all replicates on both cell lines was added to Tri Reagent containing 5 µg of tRNA carrier, and then RNA was extracted per manufacturer protocol. Two-step RT-PCR was then performed using the First-strand cDNA Synthesis M-MLV Reverse Transcriptase kit (Invitrogen) and a primer targeting EEEV 6K (**Supp. Table 1**). The E2 gene was then amplified with PfuUltra-HF per the manufacturer’s protocols using primers targeting EEEV 6K and EEEV E3 (**Supp. Table 1**). Sequencing was then carried out by Genewiz.

### Mosquito infections

All mosquito procedures were performed at the University of Texas Medical Branch in ACL3 facilities. *Aedes albopictus* (Salvador, Brazil) mosquitoes were reared and maintained at 28°C and 80% relative humidity with a 12:12 light:dark cycle with water and sucrose provided *ad libitum*.

Adult female *Ae. albopictus* mosquitoes were infected with equivalent genomes of EEEV WT, 71-77, 84-119, and 156-157 corresponding to 8.2 log_10_ WT PFU/mL in artificial bloodmeals. Engorged females were incubated at 27°C for 10 days under 12-hour light/12-hour dark circadian lighting conditions and assayed for infection by inoculation of triturated legs, saliva, and carcasses onto Vero cell monolayers and observation for cytopathic effects as previously described^5,48^.

### Mouse Studies

All animal studies were done in accordance with the recommendations in the Guide for the Care and Use of Laboratory Animals of the National Research Council. All procedures were performed at the University of Pittsburgh under protocols approved by the Institutional Animal Care and Use Committee of the University of Pittsburgh. Female 4-week-old CD-1 mice were purchased commercially (Jackson Laboratories). Virus inoculations were performed under anesthesia that was induced and maintained with isoflurane, and then mice were monitored for 14 days with approved euthanasia criteria based on weight loss and morbidity, and all efforts were made to minimize animal suffering.

For pathogenesis studies 4–6-week-old female CD-1 mice (Charles River) were infected sc. in the rear fp. with WT or mutant EEEV viruses with genome equivalents corresponding to 10^3^ WT PFU. CD-1 mice were also infected ic. with SINV/EEEV WT or mutant viruses with genome equivalents corresponding to 10^4^ WT PFU.

For VLDLR LA(1-2)-Fc decoy studies 4-6 week-old female CD-1 mice were treated with 100 µg of VLDLR LA(1-2)-Fc or LDLRAD3 D1-Fc six hours before sc. infection in the rear footpad with WT or mutant EEEV viruses with genomes equivalent to 10^3^ WT PFU. Mice were monitored for weight loss and clinical signs of illness for 14 dpi.

### Statistical Analysis

Statistical significance for all experiments was determined using Prism version 9.1 (GraphPad Software) and is indicated in each figure legend. Cell culture experiments were analyzed by one- or two-way ANOVA or repeated measures ANOVA with post-hoc tests. Mouse survival studies were performed at least twice with similar results, and the significance of survival differences was determined by the Mantel-Cox log-rank test.

### Data and material availability

All data supporting the findings of this study are available in this paper and are available upon request. This paper does not include any original code. All reagent requests and resources should be directed to the Lead Contact author. All reagents will be made available on request after completion of a Materials Transfer Agreement (MTA).

## Supplemental Information

**Supp. Fig 1.**
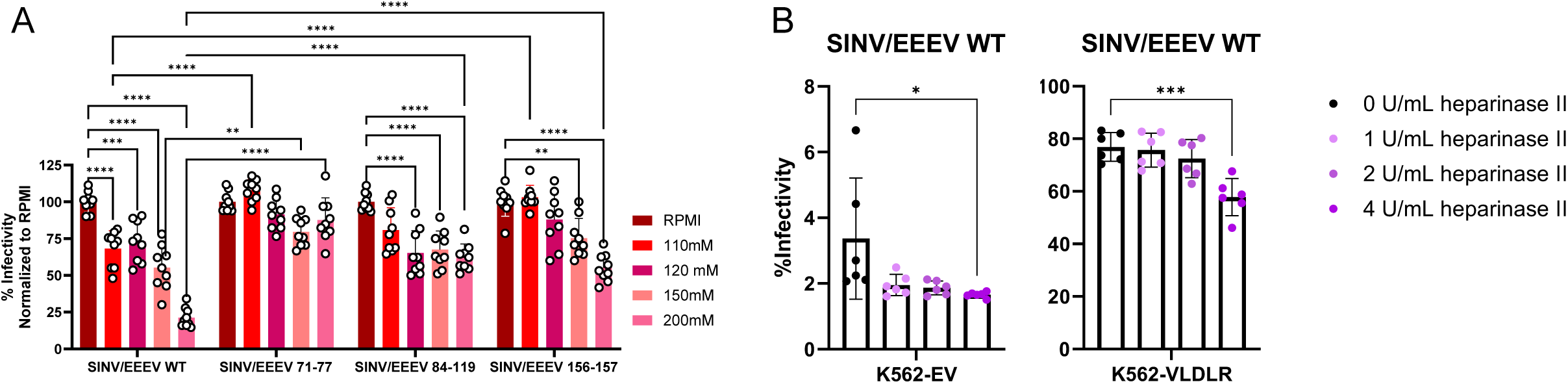
Related to Figs. 1 and 2. Impact of ionic disruption and ablation of HS-binding to SINV/EEEV WT EEEV infectivity. **A** Relative infectivity of WT or mutant SINV/EEEV diluted in RPMI supplemented with increasing NaCl concentrations on BHK-21 cells (n = 9 from 3 independent experiments). **B** Infectivity SINV/EEEV WT eGFP reporter virus of K562-EV or K562-VLDLR following digestion with heparinase II (n = 6 from 2 independent experiments). Error bars show SD. Significance was determined by (**A**) two-way ANOVA with Tukey’s post-hoc tests or (**B**) one-way ANOVA with Bonferroni’s post-hoc tests. *(p<0.05), **(p<0.01), ***(p<0.001), ****(p<0.0001).

**Supp. Fig 2.**
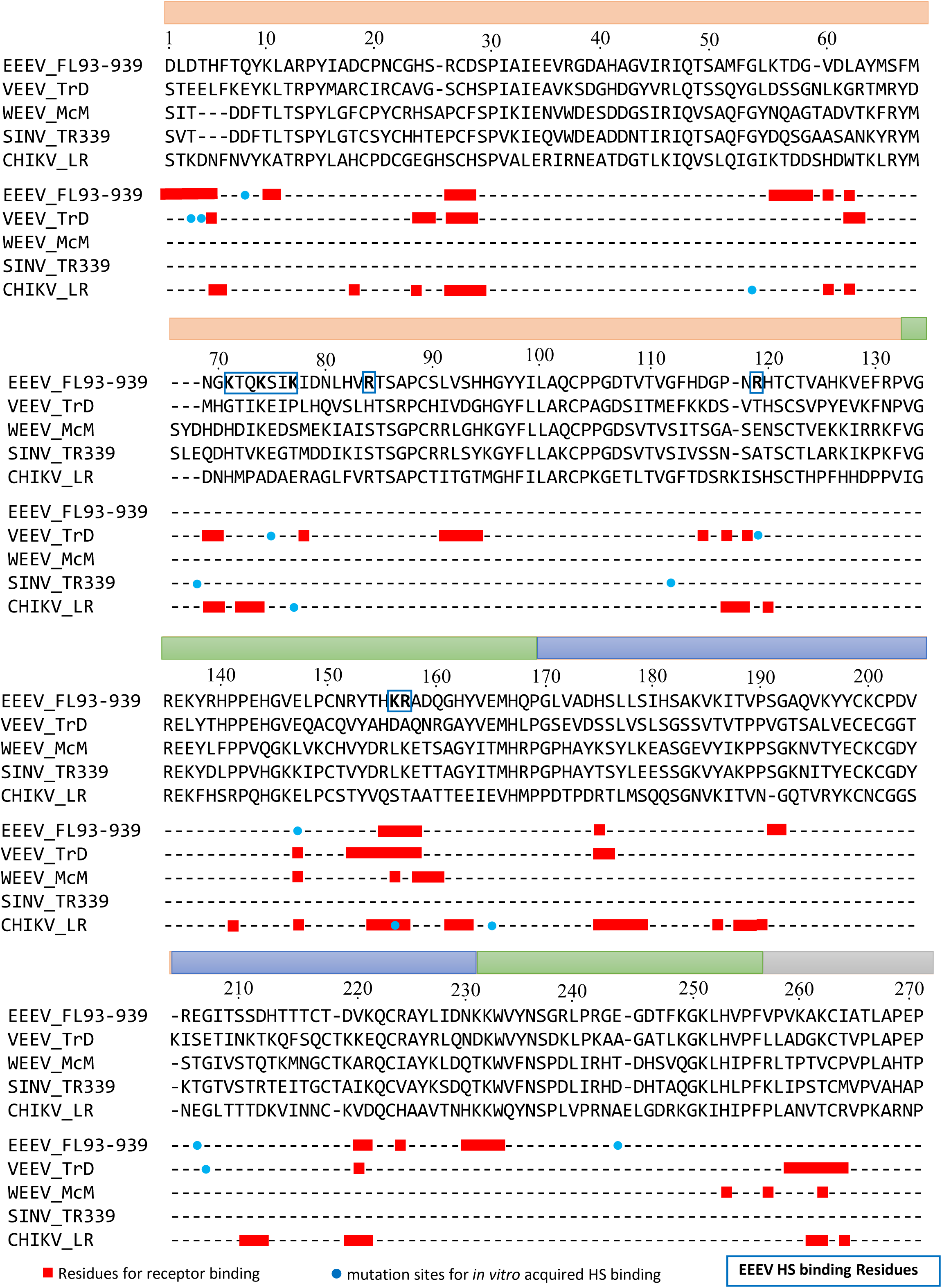
Related to Fig. 6. Sequence alignment of E2 of multiple alphaviruses indicating residues important for HS- and receptor-binding. Alphavirus strains (EEEV strain FL93-939, ABL84687; VEEV IAB strain TrD, AAC19322; WEEV strain McMillian, GQ287640; SINV strain TR339, WPT07523; CHIKV strain La Réunion, KY575571) were structurally aligned using PROMAL3D^49^ using EEEV numbering. Red boxes indicate residues that have been identified to be critical for receptor binding^11-13,29-31,33^, and blue circles represent residues where cell-culture adapted mutation to a lysine or arginine led to increased HS-binding^18,22,28^. EEEV naturally occurring HS-binding residues are boxed in blue, and residues are bolded^3,4,8^.

**Supp. Fig. 3.**
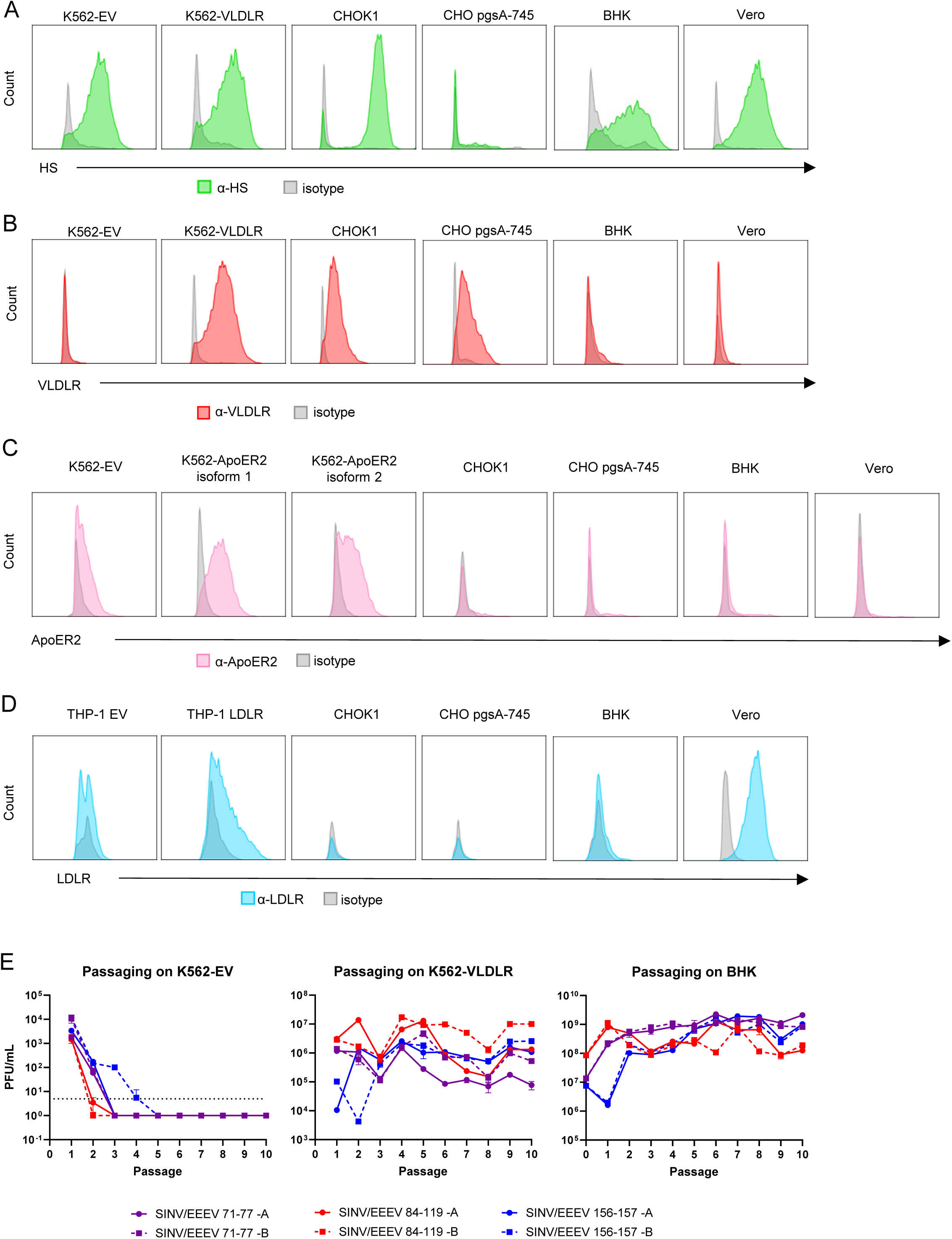
Related to Fig. 3. VLDLR and HS staining on cell lines and importance of HS for EEEV infectivity in myeloid cells. **A** Anti-HS (α-HS) cell surface staining of indicated cell types analyzed by FACS (representative staining from n = 4 from 2 independent experiments). **B** α-VLDLR cell surface staining of indicated cell types analyzed by FACS (representative staining from n = 4 from 2 independent experiments). **C** α-ApoER2 cell surface staining of indicated cell types analyzed by FACS (representative staining from n = 4 from 2 independent experiments). **D** α-LDLR cell surface staining of indicated cell types analyzed by FACS (representative staining from n = 4 from 2 independent experiments). **E** Titers of chimeric 71-77, 84-119, and 156-157 eGFP TaV during passage on K562-EV, K562-VLDLR, and BHK-21 cells. Error bars show SD.

**Supp. Fig 4.**
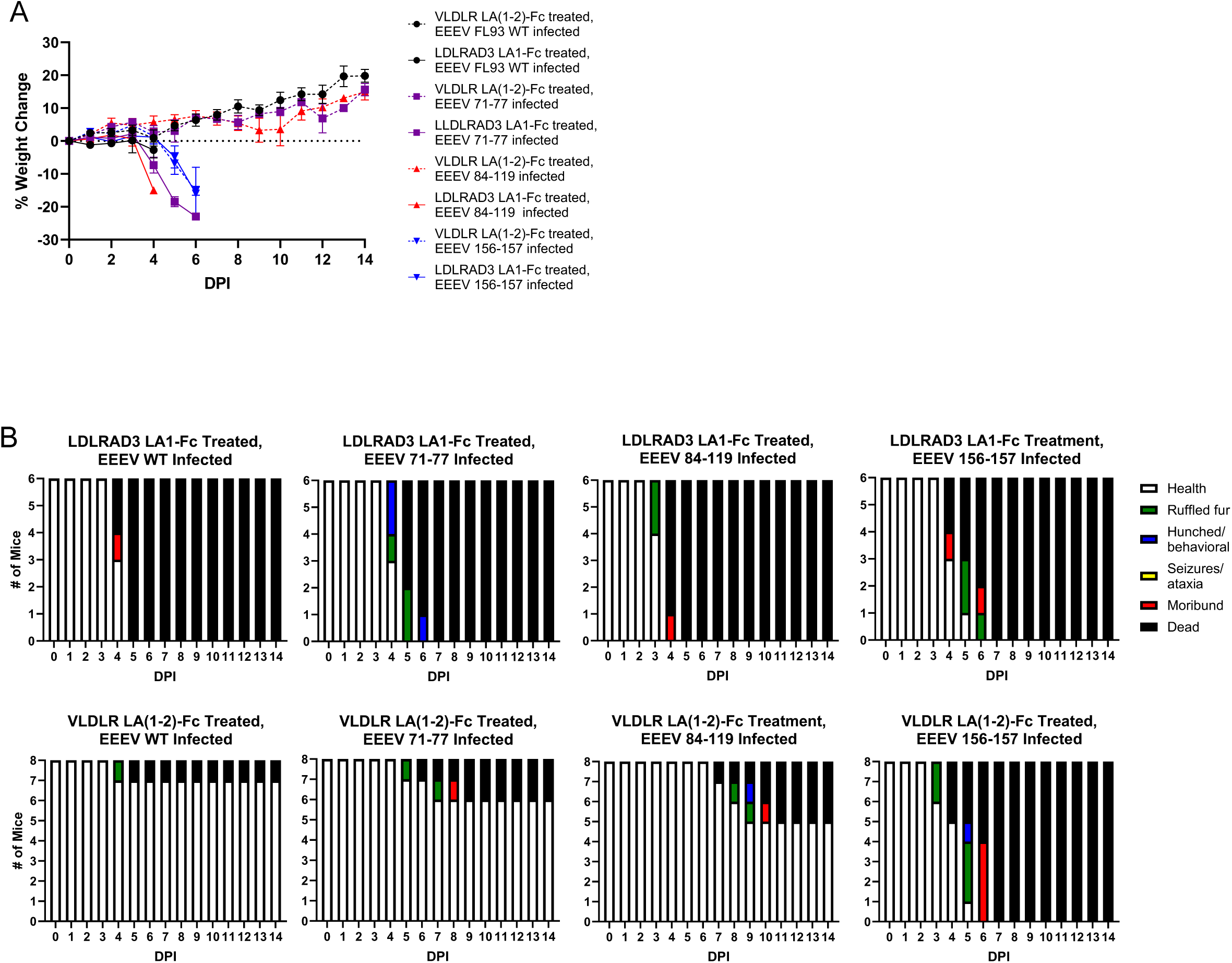
Related to Fig. 5. Weight loss and clinical signs of mice pre-treated with VLDLR-Fc decoy then infected with EEEV WT or E2 mutants. Groups of 3 or 5 mice were treated with 100 µg of VLDLR-Fc or LDLRAD3 D1-Fc as a control, 6 hours after treatment, mice were infected with were infected in the rear fp. with equal genomes of virus equivalent to 10^3^ PFU of WT (n = 6 mice for LDLRAD3 LA1-Fc and n = 8 for VLDLR LA(1-2)-Fc). **A** Weight change. **B** Clinical scores.

**Supp. Table 1.**
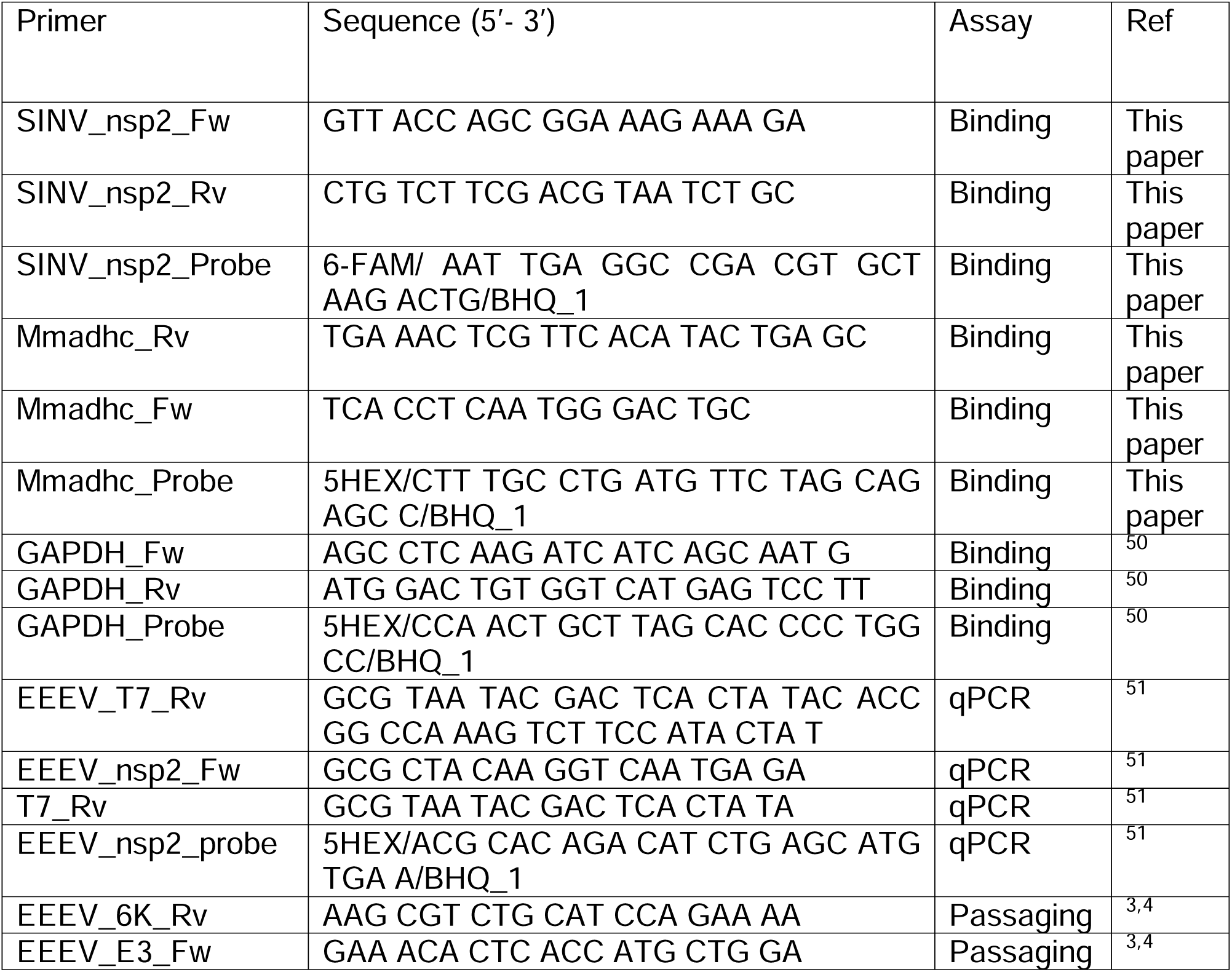
Primers used for assays.

